# Controlling pericellular oxygen tension in cell culture reveals distinct breast cancer responses to low oxygen tensions

**DOI:** 10.1101/2023.10.02.560369

**Authors:** Zachary J. Rogers, Thibault Colombani, Saad Khan, Khushbu Bhatt, Alexandra Nukovic, Guanyu Zhou, Benjamin M. Woolston, Cormac T. Taylor, Daniele M. Gilkes, Nikolai Slavov, Sidi A. Bencherif

## Abstract

Oxygen (O_2_) tension plays a key role in tissue function and pathophysiology. O_2_-controlled cell culture, in which the O_2_ concentration in an incubator’s gas phase is controlled, is an indispensable tool to study the role of O_2_ *in vivo*. For this technique, it is presumed that the incubator setpoint is equal to the O_2_ tension that cells experience (*i.e.*, pericellular O_2_). We discovered that physioxic (5% O_2_) and hypoxic (1% O_2_) setpoints regularly induce anoxic (0.0% O_2_) pericellular tensions in both adherent and suspension cell cultures. Electron transport chain inhibition ablates this effect, indicating that cellular O_2_ consumption is the driving factor. RNA-seq revealed that primary human hepatocytes cultured in physioxia experience ischemia-reperfusion injury due to anoxic exposure followed by rapid reoxygenation. To better understand the relationship between incubator gas phase and pericellular O_2_ tensions, we developed a reaction-diffusion model that predicts pericellular O_2_ tension *a priori*. This model revealed that the effect of cellular O_2_ consumption is greatest in smaller volume culture vessels (*e.g.,* 96-well plate). By controlling pericellular O_2_ tension in cell culture, we discovered that MCF7 cells have stronger glycolytic and glutamine metabolism responses in anoxia *vs.* hypoxia. MCF7 also expressed higher levels of *HIF2A*, *CD73*, *NDUFA4L2*, *etc.* and lower levels of *HIF1A*, *CA9*, *VEGFA, etc.* in response to hypoxia *vs.* anoxia. Proteomics revealed that 4T1 cells had an upregulated epithelial-to-mesenchymal transition (EMT) response and downregulated reactive oxygen species (ROS) management, glycolysis, and fatty acid metabolism pathways in hypoxia *vs.* anoxia. Collectively, these results reveal that breast cancer cells respond non-monotonically to low O_2_, suggesting that anoxic cell culture is not suitable to model hypoxia. We demonstrate that controlling atmospheric O_2_ tension in cell culture incubators is insufficient to control O_2_ in cell culture and introduce the concept of *pericellular O_2_-controlled cell culture*.

## Introduction

A cornerstone of biological research, cell culture aims to grow cells in conditions that simulate their native environment as closely as possible. Cell culture models serve as a tool for testing biological hypotheses before validating *in vivo*. Healthy and diseased tissues are isolated from patients and studied *in vitro*. In fact, cell culture techniques are used throughout the process of drug development to make “go/no-go” decisions^1, 2^ and to manufacture adoptive cell therapies and regenerative medicines.^3, 4^ Recent advances in this practice include scaffolds that mimic the extracellular matrix^5–7^, self-assembly of pluripotent stem cells to form brain organoids^8^, patient-derived organoids that capture tumor heterogeneity in patients and predict therapeutic responses^9, 10^, *etc.* Yet, one key discrepancy between *in vitro* and *in vivo* remain: the “normoxic” oxygen (O_2_) tension in cell culture (141 mmHg) is dramatically higher than the O_2_ tension of human tissues (3– 100 mmHg) (1% O_2_ = 7.7 mmHg O_2_ at sea level)).^11, 12^

Supraphysiological O_2_ concentrations (hyperoxia) lead to excessive reactive oxygen species (ROS) production, resulting in cellular damage and dysregulated signaling.^13, 14^ It is therefore not surprising that cells grown in physioxia experience less oxidative stress.^15, 16^ Furthermore, hyperoxia degrades proteins containing a specific iron-sulfur cluster, disrupting diphthamide synthesis, purine metabolism, nucleotide excision repair, and electron transport chain (ETC) function.^17^ Many O_2_-dependent enzymes require iron and copper metal cofactors, which are susceptible to oxidation in hyperoxia.^12, 18^ Studies culturing cells in normoxia *vs.* physioxia have found aberrant T-cell activation^19^, fibroblast senescence^20^ and mutation frequency^21^, chondrocyte differentiation^22^, *etc.* in normoxia. However, the full impact of culturing cells in normoxia will remain unknown until physioxia becomes common practice.

To address these concerns, tools to perform physiological cell culture have been developed and are commercially available. These products, including portable chambers, tri-gas incubators, and hypoxic workstations, consist of chambers that control O_2_ in the atmosphere of cultured cells by adding compressed nitrogen. However, widespread adoption has been hampered by cost, laboratory space requirements, technical challenges, and rapid reoxygenation of cultures.^23, 24^ Reoxygenation, which occurs when cell cultures are removed from portable chambers or tri-gas incubators and exposed to normoxia, makes it difficult to recapitulate physiological O_2_ tensions.^18, 25^ One major application for these products are hypoxic cell culture models, conducted at 0.5 – 1% O_2_.^26–30^ Hypoxia occurs in both physiological (*e.g.,* placenta, renal medulla, intestinal mucosa, germinal centers, bone marrow) and pathophysiological (*e.g.,* infection, inflammation, solid tumors, ischemia) contexts; and is therefore an active area of research.^31, 32^ Hypoxic cell culture was instrumental in the discovery of the prolyl hydroxylase (PHD) / hypoxia-inducible factor (HIF) axis, the mechanism by which cells sense and respond to low O_2_.^33, 34^ O_2_-controlled cell culture is also performed to mimic physioxia, typically at 5% O_2_.^15, 24^

In O_2_-controlled cell culture, it is generally presumed that the atmospheric O_2_ tension within incubators is equal to the pericellular O_2_ tension, the O_2_ concentration that cells experience. The pericellular O_2_ tension is dependent on several rates: the O_2_ diffusion within the cell culture media, O_2_ transfer at the gas–media interface, and the cellular O_2_ consumption. Gas–media O_2_ transfer is the limiting rate.^35, 36^ Culture vessel geometry, medium volume, and surface area also influence diffusion times. These parameters vary greatly based on user preference and experimental design, yet are not reported. Experimental studies measuring pericellular O_2_ tension indicate that confluent normoxic cultures can induce hypoxia.^37^ However, the impact of O_2_ consumption rates in lower O_2_ tensions is unclear, since consumption decreases as O_2_ becomes limiting.^38^ We set out to assess how key cell culture parameters (*i.e.*, cell type, cell density, medium volume, and culture vessel geometry) influence the relationship between atmospheric and pericellular O_2_ tensions in O_2_-controlled cell culture models. After discovering that pericellular O_2_ tension is vastly different from atmospheric O_2_ tension, we explored how controlling pericellular O_2_ tension could be used as a novel tool to study breast cancer cell responses in low O_2_.

## Results

### 1% O_2_ media conditioning can take over 5 days and is reoxygenated within minutes

For O_2_-controlled cell culture experiments, media is typically conditioned to the desired O_2_ tension and added to the cells at the start of the experiment. This procedure ensures that cells experience the desired O_2_ tension immediately. We investigated how long it would take to condition 25–500 mL of media for hypoxic (1% O_2_) experiments, since conditioning times are not reported.^26, 28, 39^ The required time was far longer than anticipated: over 1 day for 25 mL (upright T75 flask) and over 5 days for 500 mL (**Figure 1A**). The type of culture vessel or tube used did not change conditioning time for 25 or 50 mL of media. To understand media reoxygenation kinetics, 500 mL of 1% O_2_ medium was removed from a tri-gas incubator and aliquoted into different culture vessels containing O_2_ sensors. By the time the media reached the culture vessels, the O_2_ concentration was >6% O_2_ (**Figure 1B**). Depending on the surface area of the medium in different culture vessels, the medium reached 10% O_2_ within seconds to 10 minutes. These results indicate that 1% O_2_ media conditioning requires surprisingly long periods – more than 5 days for large volumes. Furthermore, 1% O_2_ media is rapidly reoxygenated when removed from controlled O_2_ atmospheric environments, indicating that portable chambers and tri-gas incubators are not suitable to condition media.

**Figure 1:**
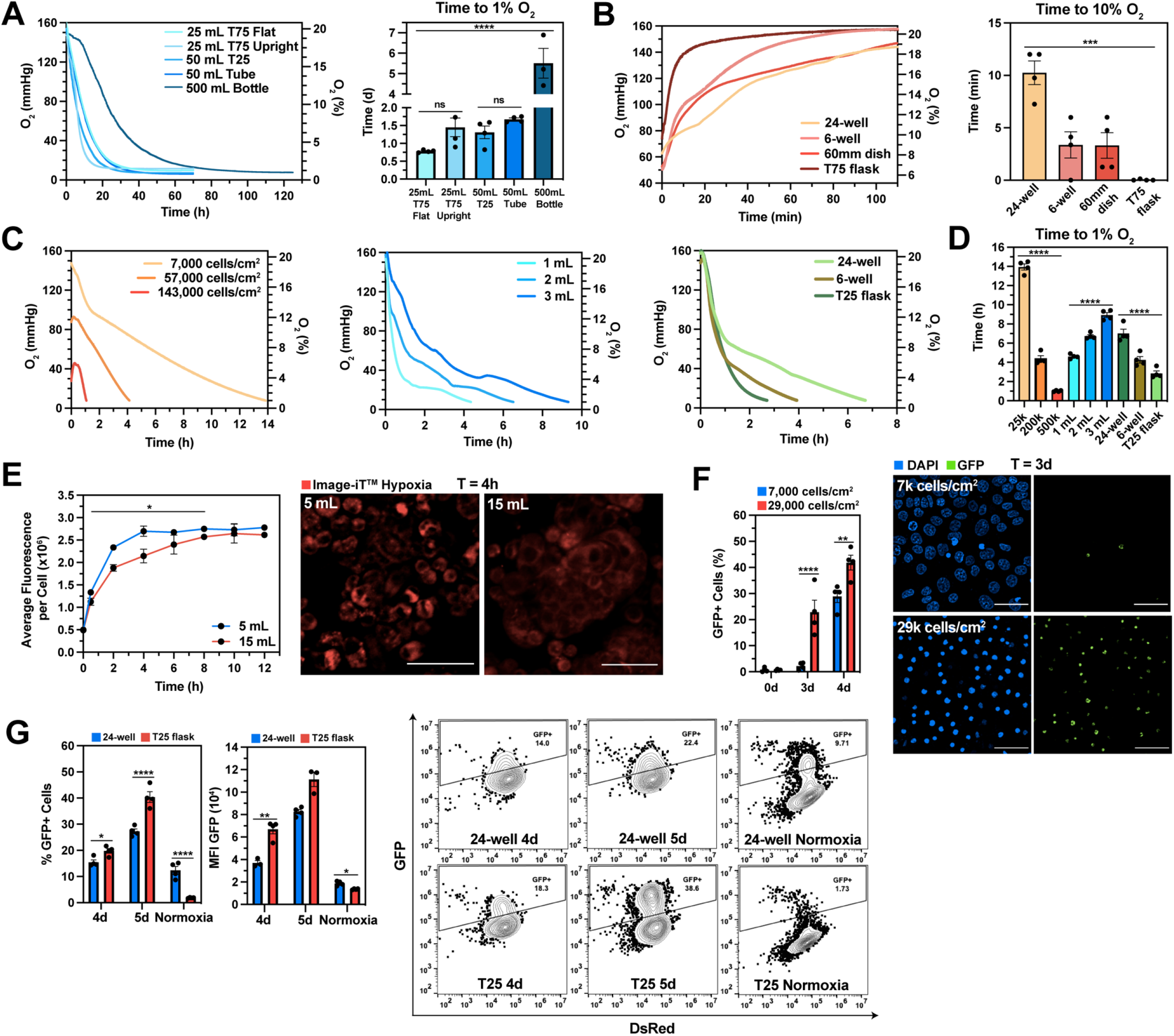
C**e**ll **culture parameters influence HIF stabilization kinetics in hypoxic (1% O_2_) culture. (A)** O_2_ kinetics (left) and time to 1% O_2_ (right) of normoxic media (EMEM + 10% FBS + 1% P/S) placed in a 1% O_2_ incubator. **(B)** O_2_ kinetics (left) and time to 10% O_2_ (right) of 1% O_2_ media placed into different culture vessels under normoxia. **(C)** O_2_ kinetics of MCF7 cultures with varying cell densities (left), medium volumes (middle), and culture vessel type (right) placed in a 1% O_2_ incubator. **(D)** Time to 1% O_2_ from (C). **(E)** Average fluorescence (Image-iT^TM^ Hypoxia) per cell (left) and representative confocal images at 4 hours (right) for MCF7 cultures (60mm dish) with 5 or 15 mL of media placed in a 1% O_2_ chamber. Red = Image-iT^TM^ Hypoxia as an indicator of cellular hypoxia. **(F)** Percentage of GFP+ cells (left) and representative confocal images at 3 days (right) from MCF7 HIF reporter cells cultured at 7,000 or 29,000 cells/cm^2^ for 4 days in a 1% O_2_ incubator. Blue = nuclei stained with DAPI, Green = GFP+ (HIF+) cells. **(G)** Percentage of GFP+ cells (left), GFP mean fluorescence intensity (MFI) (middle) and representative contour plots with outliers (right) for MDA-MB-231 HIF reporter cells cultured in a 24-well plate or T25 flask for up to 5 days in a 1% O_2_ incubator. Data were analyzed with ANOVA and Geisser-Greenhouse (E) or Bonferroni (F-G) corrections. N = 3 – 4 biological replicates per condition.

### Cell density, medium volume, and culture vessel type influence HIF stabilization kinetics

We investigated how cell density, medium volume, and culture vessel type influence the time it takes normoxic cultures to reach 1% O_2_. Non-invasive optical sensor spots were used to measure pericellular O_2_ tension.^40, 41^ For MCF7 breast cancer cultures, all three parameters influenced the time to 1% O_2_, ranging from 1–14 hours (**Figure 1C–D**). Cell density had the largest effect, suggesting that cellular O_2_ consumption plays a key role in the induction of hypoxia *in vitro*.

Next, we explored whether medium volume influenced cellular hypoxia kinetics. MCF7 cultures in 60mm dishes containing either 5 mL or 15 mL of media were placed inside a 1% O_2_ incubator. Cellular hypoxia was evaluated using a hypoxia-responsive fluorescent dye (Image-iT^TM^ Hypoxia) for 12h. As expected, the cells in the 5 mL condition reached a maximum cellular fluorescence sooner than the 15 mL cultures: 4 hours *vs.* 10 hours (**Figure 1E, Figure S1**).

To understand if cell density or culture vessel type influenced HIF stabilization in 1% O_2_ culture, MCF7 and MDA-MB-231 HIF reporter cell lines^28^ were cultured at two cell densities for 4 days and the percentage of GFP-positive cells was determined using fluorescent microscopy. After 3 days, 2.2% of cells plated at the lower cell density were GFP-positive whereas 22.9% of cells plated at a higher density were GFP-positive (**Figure 1F, Figure S2**). We then tested whether different culture vessel types would induce a similar effect for MDA-MB-231 reporter cells. Consistent with our findings in **Figure 1C**, T25 flask cultures had a higher percentage of GFP-positive cells after 4 and 5 days compared to 24-well cultures (**Figure 1G**). Collectively, we have demonstrated that cell density, medium volume, and culture vessel type, parameters that vary between experiments and are not reported, greatly influence cellular hypoxia and HIF stabilization kinetics. These results suggest that consistent HIF stabilization kinetics in 1% O_2_ culture can only be obtained by using a workstation and conditioned media.

### Cellular O_2_ consumption can induce anoxia in both 5% and 1% O_2_ culture

After discovering that cellular O_2_ consumption drives the induction of hypoxia in 1% O_2_ incubators, we hypothesized that it would also influence the pericellular O_2_ tension. To test this, sub-confluent (21,000 cells/cm^2^) MCF7, MDA-MB-231, and primary human mammary epithelial cells were cultured in physioxic (5% O_2_) and hypoxic (1% O_2_) conditions for 72 hours and the pericellular O_2_ concentration was measured. Cell-free media O_2_ tensions matched the incubator setpoints, indicating that the O_2_ sensor spots were accurately recording and the incubator O_2_ sensors were calibrated (**Figure 2A–B**).

**Figure 2:**
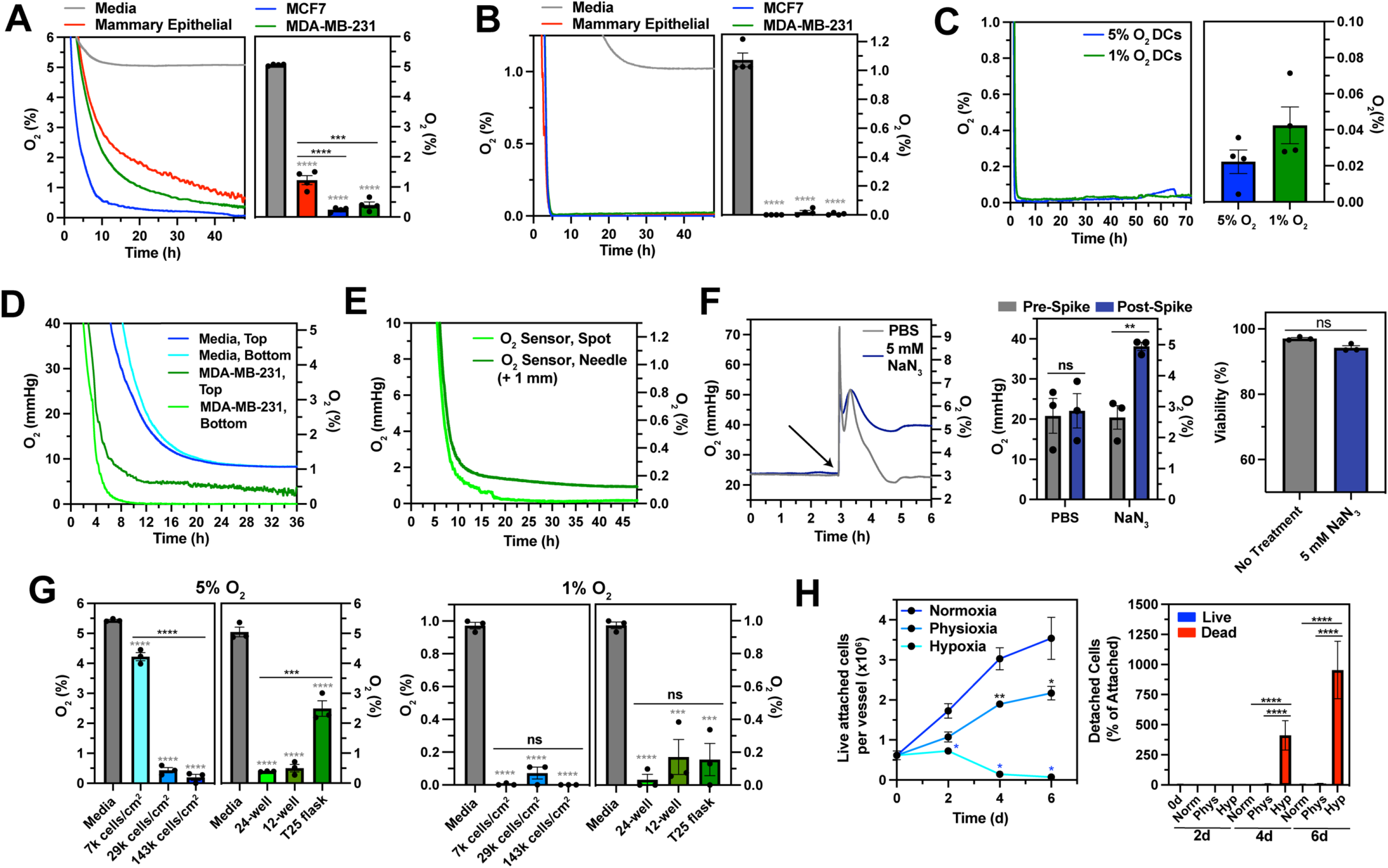
C**e**llular **O_2_ consumption regularly induces anoxia (0.0% O_2_) in both physioxic (5% O_2_) and hypoxic (1% O_2_) culture. (A–B)** O_2_ profiles (left) and average O_2_ tension values (right) of media, human mammary epithelial, MCF7, or MDA-MB-231 cultures seeded onto a 24-well plate in a 5% O_2_ (A) or 1% O_2_ (B) incubator for 72h. **(C)** O_2_ profiles (left) and average O_2_ tension values (right) of human dendritic cell (DC) 96- well cultures in a 5% or 1% O_2_ incubator for 72h. **(D)** O_2_ kinetics of media (green) or MDA- MBA-231 cultures (blue) of the top (dark) or bottom (light) of the well in a 1% O_2_ incubator. **(E)** O_2_ profiles of MDA-MB-231 cultures measured at the bottom of the well (spot) or 1mm above the bottom of the well (needle) in a 1% O_2_ incubator. **(F)** Effect of sodium azide (NaN3) on MCF7 pericellular O_2_ tension and cytotoxicity. O_2_ profiles (left) and average O_2_ tension values (middle) of MCF7 cultures spiked with either 5 mM NaN3 or PBS in a 1% O_2_ incubator. MCF7 cell viability after ± incubation with 5 mM NaN3 for 6 hours in a 1% O_2_ incubator (right). **(G)** Average O_2_ tension values of media and MCF7 cultures with different cell densities and culture vessel types in a 5% O_2_ (left) or 1% O_2_ incubator (right) for 72h. **(H)** Live, attached (left) and detached (right) cell counts for MCF7 cultures in a 18.6%, 5% or 1% O_2_ incubator for 6 days. Data were analyzed with two-tailed t test (F) or ANOVA and Dunnett’s (A-B) or Tukey’s (C, F, G, H) corrections. N = 3 – 4 biological replicates per condition. Colored * indicates comparison to the control.

In physioxia, the pericellular O_2_ was strikingly low – below 1.5% O_2_ for all three lines, and MCF7 cultures were anoxic (anoxia defined as <0.5% O_2_^42^). In hypoxia, the pericellular O_2_ of all cultures became anoxic (0.0% O_2_) within 5 hours. Next, we investigated whether this effect also occurred in suspension culture. Activated human dendritic cells (DCs) were cultured in physioxia or hypoxia for 72 hours. Both cultures were immediately and sustainably anoxic (0.0% O_2_) (**Figure 2C**).

If cellular O_2_ consumption did indeed affect pericellular O_2_ tension, there would be an axial O_2_ gradient in these cultures. To test this hypothesis, O_2_ tension at the media-gas interface of MDA-MB-231 cultured at 1% O_2_ was measured using needle O_2_ microsensors (top) and pericellular O_2_ tensions were measured using sensor spots (bottom). **Figure 2D** illustrates that in these cultures, there is a gradient of 0.4 – 0.0% O_2_ from the top to the bottom of the well after 36 hours. In wells without cells, both the top and bottom of the well reached 1% O_2_ as expected. To validate O_2_ sensor spot readings, needle microsensors were placed 1mm above sensor spots in 1% O_2_ MDA-MB-231 cultures using a micromanipulator. Sensor spots recorded 0.0% O_2_ and needles recorded 0.1% O_2_, indicating that the two probes were in good agreement (**Figure 2E**).

To further validate the role of cellular O_2_ consumption in pericellular O_2_ tension, we tested whether inhibiting oxidative phosphorylation would ablate axial gradients. Upon addition of sodium azide (NaN3, complex IV inhibitor), the pericellular O_2_ tension of MCF7 cultures rapidly rose from 3% O_2_ to the incubator setpoint of 5% O_2_, whereas PBS (vehicle) spiked cultures returned to 3% O_2_ with continued incubation (**Figure 2F, left panel**). To confirm that this effect was not due to NaN3 cytotoxicity, we confirmed that there was no significant change in MCF7 cell viability when incubated with NaN3 at 5% O_2_ for 6h (**Figure 2F, right panel**).

We next explored how cell density and culture vessel type influenced the gradient between atmospheric and pericellular O_2_ tensions. In physioxic MCF7 cultures, cell densities of 7,000, 29,000, and 143,000 cells/cm^2^ in a 12-well plate induced pericellular O_2_ tensions of 4.2%, 0.5%, and 0.1% O_2_, respectively. Different culture vessels (24-well plate, 12-well plate, and T25 flask) also influenced MCF7 tensions, ranging from 0.4 – 2.5% O_2_ (**Figure 2G, left panel**). In hypoxia, neither cell density nor culture vessel type change pericellular tensions in MCF7 cultures: all were anoxic (<0.2% O_2_) (**Figure 2G, right panel**).

After determining that all hypoxic MCF7 cultures tested were anoxic, we sought to understand how pericellular anoxia affected cell viability. MCF7 cells were cultured in normoxia, physioxia, or hypoxia for 6 days and cell proliferation and viability were evaluated. Cells cultured in normoxia and physioxia proliferated throughout the 6 day period (**Figure 2H, left panel**). However, cells cultured in hypoxia did not proliferate, and the majority were dead and detached after 4 days (**Figure 2H, right panel**).

Collectively, these experiments show that cellular O_2_ consumption drives pericellular O_2_ far below the incubator setpoint, inducing anoxia in both physioxic and hypoxic MCF7, MDA-MBA-231, and human DC cultures. Furthermore, in physioxic MCF7 culture, pericellular O_2_ tension is highly dependent on cell culture parameters, ranging from 0.1 – 4.2% O_2_.

### Setting the incubator to physioxia mimics ischemia-reperfusion injury in hepatocyte culture

We next explored how the difference between the incubator setpoint and pericellular O_2_ tension can impact the physiological relevance of cell culture models. Because of their high O_2_ consumption rate^24, 43^ and widespread use as an *in vitro* drug metabolism model^44^, primary human hepatocytes were used for these studies. Hepatocytes were seeded and cultured in either normoxic (18.6% O_2_) or physioxic (6% O_2_^24^) conditions for 36 hours (**Figure 3A**).

**Figure 3:**
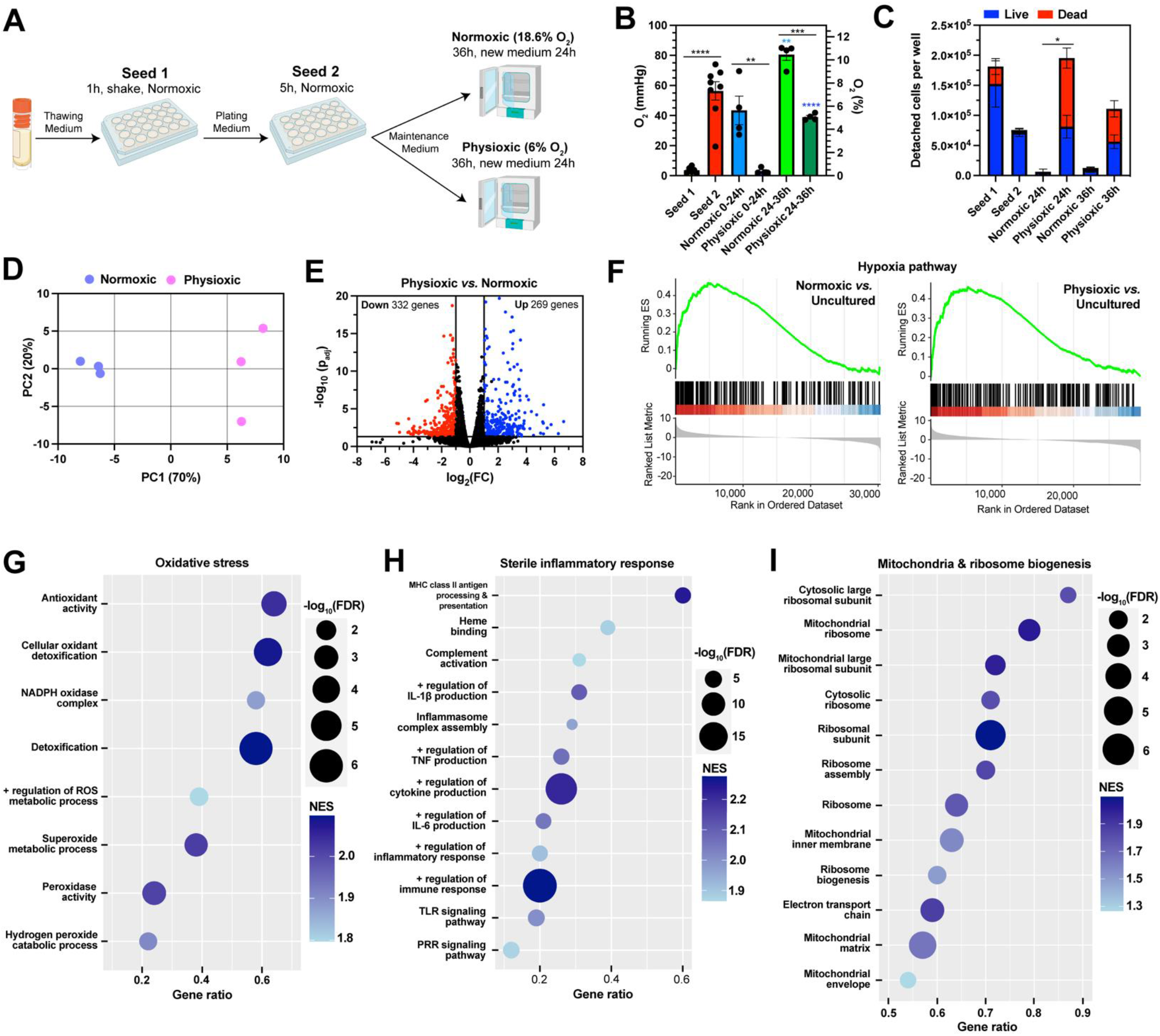
S**e**tting **the incubator to physiological O_2_ conditions mimics ischemia-reperfusion injury in human hepatocyte culture. (A)** Schematic of the primary human hepatocyte culture. RNA-seq was performed on uncultured hepatocytes and cells cultured in normoxic (18.6% O_2_) or physioxic (6% O_2_) incubator after 36h. **(B–C)** Average O_2_ tension values (B) and detached cells per well (C) during each step of the culturing process. **(D)** Principal component analysis (PCA) of the RNA-seq results for normoxic and physioxic cultured hepatocytes. **(E)** Volcano plot indicating upregulated (blue) and downregulated (red) genes for physioxic *vs.* normoxic cultured hepatocytes (padj < 0.05 and |log2FC| ≥ 1). **(F)** Hypoxia gene set enrichment analysis (GSEA) from the Hallmark database for normoxic *vs.* uncultured (left) and physioxic vs. uncultured (right). **(G–I)** Enriched pathways from the Gene Ontology (GO) database for physioxic *vs.* normoxic samples associated with oxidative stress **(G)**, sterile inflammatory response **(H)** and mitochondrial and ribosomal biogenesis **(I)**. Data were analyzed with ANOVA and Tukey’s correction (B-C). N = 3 biological samples per condition for RNA seq analysis.

During the first seeding step, which was conducted in normoxia, hepatocytes were anoxic (0.5% O_2_) (**Figure 3B**). During the first 24 hours of culture, hepatocytes cultured in normoxia were physioxic (5.6% O_2_) and hepatocytes cultured in physioxia were anoxic (0.4% O_2_). Unexpectedly, when media was exchanged after 24 hours, both types of cultures underwent reoxygenation for the duration of the experiment. **Figure 3C** illustrates that the pericellular anoxia experienced by hepatocytes cultured in physioxia increased the number of detached cells after 24 hours, the majority of which were dead.

To investigate how the pericellular O_2_ tension influenced hepatocyte physiology, we assessed gene expression by RNA-seq of uncultured hepatocytes, and hepatocytes cultured in normoxia or physioxia after 36h. Principal component analysis (PCA) of the transcriptome shows clustering of replicates by O_2_ tension, as expected (**Figure 3D**). Differential gene expression analysis (physioxic *vs.* normoxic) found 269 upregulated and 332 downregulated genes (**Figure 3E**). Gene set enrichment analysis (GSEA) revealed an upregulation in hypoxia-associated genes in normoxic and physioxic cultured cells compared to uncultured hepatocytes (**Figure 3F**), indicating that the 1-hour anoxic seeding step induced a hypoxic response. In addition, GSEA suggests that hepatocytes cultured in physioxia mounted an oxidative stress response (**Figure 3G**). Corroborating the higher cell death and detachment, physioxic cultured hepatocytes also had an upregulation in IL-1β production, TNF production, TLR signaling, and PRR signaling pathways, indicating a sterile inflammatory response (**Figure 3H**). Lastly, hepatocytes in physioxia had upregulated mitochondrial and ribosomal biogenesis pathways (**Figure 3I**). Taken together, these results indicate that setting the incubator to physiological O_2_ conditions mimics ischemia-reperfusion injury in hepatocytes due to their cellular O_2_ consumption.

### Developing a reaction-diffusion model to predict pericellular O_2_ tension in cell cultures

Measuring pericellular O_2_ tension for every O_2_-controlled cell culture experiment would be cumbersome and expensive. We hypothesized that a computational model could predict pericellular tension *a priori*, given cell density, O_2_ consumption rate, culture vessel type and medium volume. Such a tool would reduce the need for experimental measurements.

We first examined whether the unsteady state diffusion equation^35, 36^ could describe O_2_ transfer kinetics between cell culture medium and incubator gas phases. Coefficient of determination (R^2^) values suggested that experimental and numerical values that describe the O_2_ transfer between normoxic media and 1% O_2_ gas phase in different culture vessels were in good agreement (**Figure 4A**). The diffusion model also predicted equilibration of normoxic media in a 5% O_2_ incubator (**Figure S3A**). In addition, the diffusion model’s analytical solution was also comparable to numerical solutions for different culture vessel types (**Figure S3B**).

**Figure 4:**
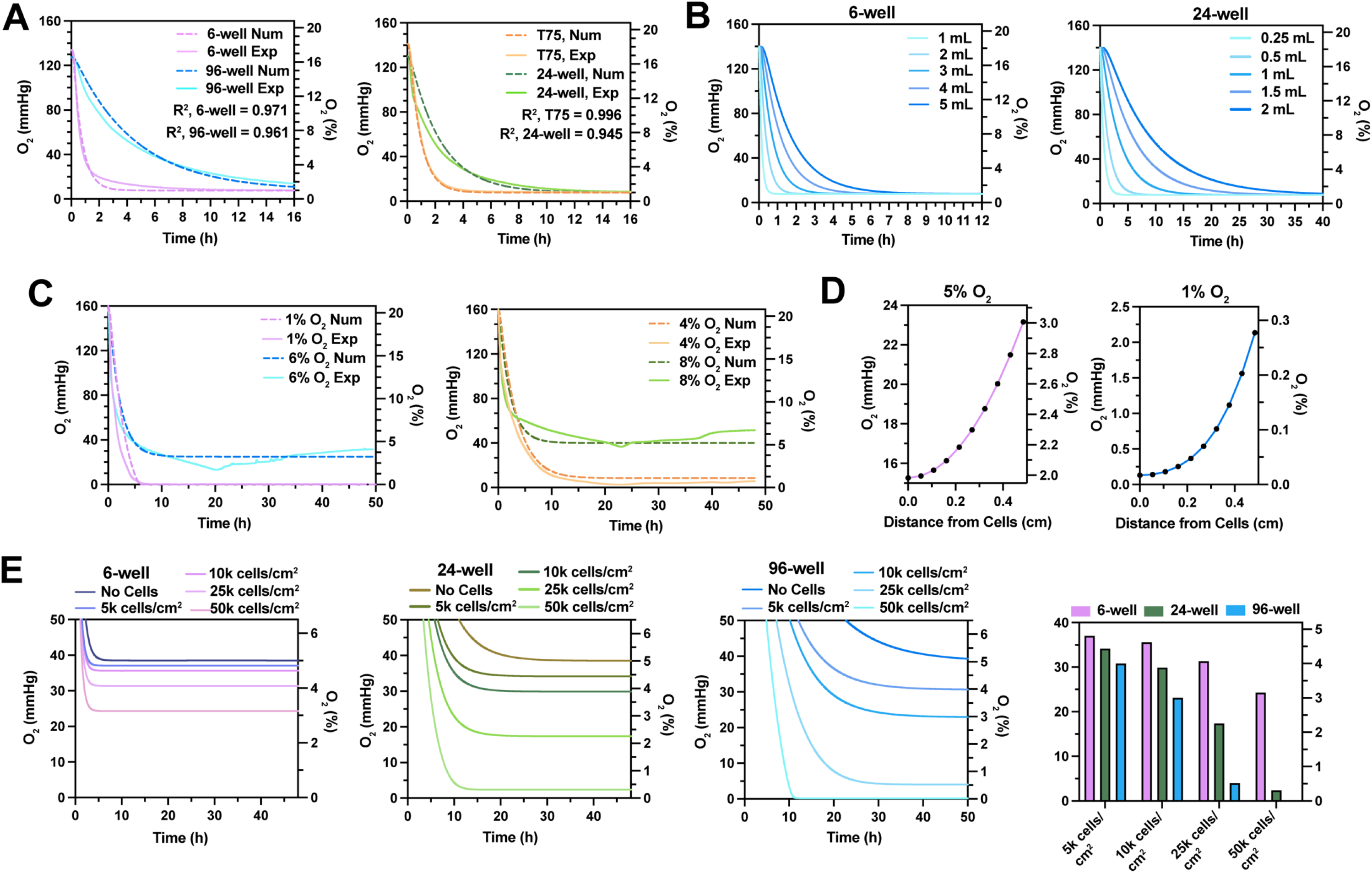
Developing a reaction-diffusion model to predict pericellular O_2_ tension in cell cultures. (A) Numerical (num) (dashed) and experimental (exp) (solid) O_2_ kinetics of normoxic media (EMEM + 10% FBS + 1% P/S) in different culture vessels placed in a 1% O_2_ incubator. (B) Diffusion model predictions for different volumes of normoxic media in a 6-well plate (left) or a 24-well plate (right) placed in a 1% O_2_ incubator. (C) Numerical and experimental O_2_ kinetics of MDA-MB-231 cultures seeded in a 24-well plate placed in a 1%, 4%, 6%, or 8% O_2_ incubator for 48h. (D) Reaction-diffusion model predictions of the gradient within MDA-MB-231 cultures seeded in a 24-well plate in a 5% O_2_ (left) or 1% O_2_ (right) incubator. (E) Reaction-diffusion model predictions of MDA-MB-231 cells cultured at different cell densities and culture vessel types in a 5% O_2_ incubator. N = 4 biological replicates per condition for experimental data.

After validating the diffusion model, we applied it to investigate the dependency of O_2_ transfer kinetics on medium volume in a 24-well plate and 6-well plate. **Figure 4B** shows that although commonly used medium volumes for 24-well and 6-well plates modestly impact the time to 1% O_2_, there is a substantial difference in kinetics between 24-well and 6-well plate wells. 6-well plate volumes of media require 0.5–12 hours to reach 1% O_2_, whereas 24-well plate requires 2.5–40 hours. Importantly, the rate of equilibration for the same medium volumes (1 mL and 2 mL) is substantially lower in 24-well plate than 6-well plate wells. These findings indicate that the gas phase–pericellular O_2_ differential increases as culture vessel surface area decreases (**Figure 2G**) because of a decrease in O_2_ transfer rates.

Next, we developed a reaction-diffusion model to describe pericellular O_2_ tension in cell cultures within O_2_-controlled environments. Michaelis–Menten kinetics were used to model cellular O_2_ consumption.^38^ This model predicts pericellular O_2_ tension values for MDA-MB-231 cultured at 1%, 4%, 6%, and 8% O_2_ with reasonable Michael–Menten parameters (V_max_ = 450 amol cell^-1^ sec^-1^ and Km = 1 µM)^38^ (**Figure 4C**). For further validation, we tested whether the model predicted axial O_2_ gradients like those experimentally determined in **Figure 2D**. The model predicts gradients of 2.0 – 3.0% O_2_ and 0.0 – 0.3% O_2_ for 30,000 cells/cm^2^ MDA-MB-231 cultures in physioxia and hypoxia, respectively (**Figure 4D**), in good agreement with experimental data.

Using the reaction-diffusion model, we next examined the influence of cell density on pericellular O_2_ tension in different culture vessel types in physioxia. The model predicts that the highest cell density will have a modest influence on pericellular tension in 6-well plate cultures (3.2% O_2_), but will induce anoxia in 24-well (0.4% O_2_) and 96-well plate cultures (0.0% O_2_) (**Figure 4E**). Lastly, we investigated the influence of cellular O_2_ consumption (V_max_) in hypoxia (**Figure S3C**). Both cell density and O_2_ consumption predictions suggest that the smaller the culture vessel surface area, the greater the impact of cellular O_2_ consumption on pericellular O_2_ tension. This effect is due to the decreasing media surface area to height as culture vessel size decreases (**Figure S4D**). Taken together, we establish that a reaction-diffusion model can predict pericellular O_2_ tension in O_2_-controlled cell culture. Furthermore, using the model, we discovered that the effect of cellular consumption on pericellular O_2_ tension is highly dependent on the culture vessel type, increasing as culture vessel size decreases.

### Pericellular anoxia induces stronger metabolic reprogramming compared to pericellular hypoxia in MCF7 cells

The studies presented thus far demonstrate that standard hypoxic cell culture (1% O_2_) routinely induces anoxia due to cellular O_2_ consumption. Because anoxia is not physiologically relevant *in vivo*, we asked whether anoxia is suitable to model hypoxia. To explore this concept, we controlled pericellular O_2_ tension to investigate cancer cell responses to pericellular hypoxia (1–2% O_2_) *vs.* pericellular anoxia (0% O_2_) .

First, we examined MCF7 metabolic reprogramming in response to different pericellular O_2_ tensions. Expected metabolic changes in response to hypoxia include an increase in (i) glucose consumption due to increased uptake and glycolytic flux, (ii) extracellular lactate from decreased TCA cycle flux and increased lactate transport, (iii) glutamine uptake to replenish TCA cycle intermediates for lipid metabolism, and (iv) extracellular glutamate secretion, which promotes cancer cell proliferation.^39, 42, 45–48^ MCF7 cells were cultured for 72 hours in 18.6%, 4%, and 1% O_2_ incubators, resulting in supraphysiologic (10.8% O_2_), hypoxic (1.2% O_2_), and anoxic (0.0% O_2_) pericellular tensions, respectively (**Figure 5A**). Consumption (glucose and glutamine) and production (lactate and glutamate) rates trended higher with decreasing pericellular O_2_ tension over the 72 hour time course (**Figure 5B – 5E**). For example, lactate and glutamate production was 2.2- and 1.6-fold higher for cells in anoxia compared to hypoxia, respectively. Only the anoxic MCF7 cultures exhibited increased rates after 24 hours, whereas hypoxic and normoxic cultures maintained constant rates. These findings suggest that for MCF7 cells, anoxia induces a stronger metabolic reprogramming response than hypoxia does.

**Figure 5:**
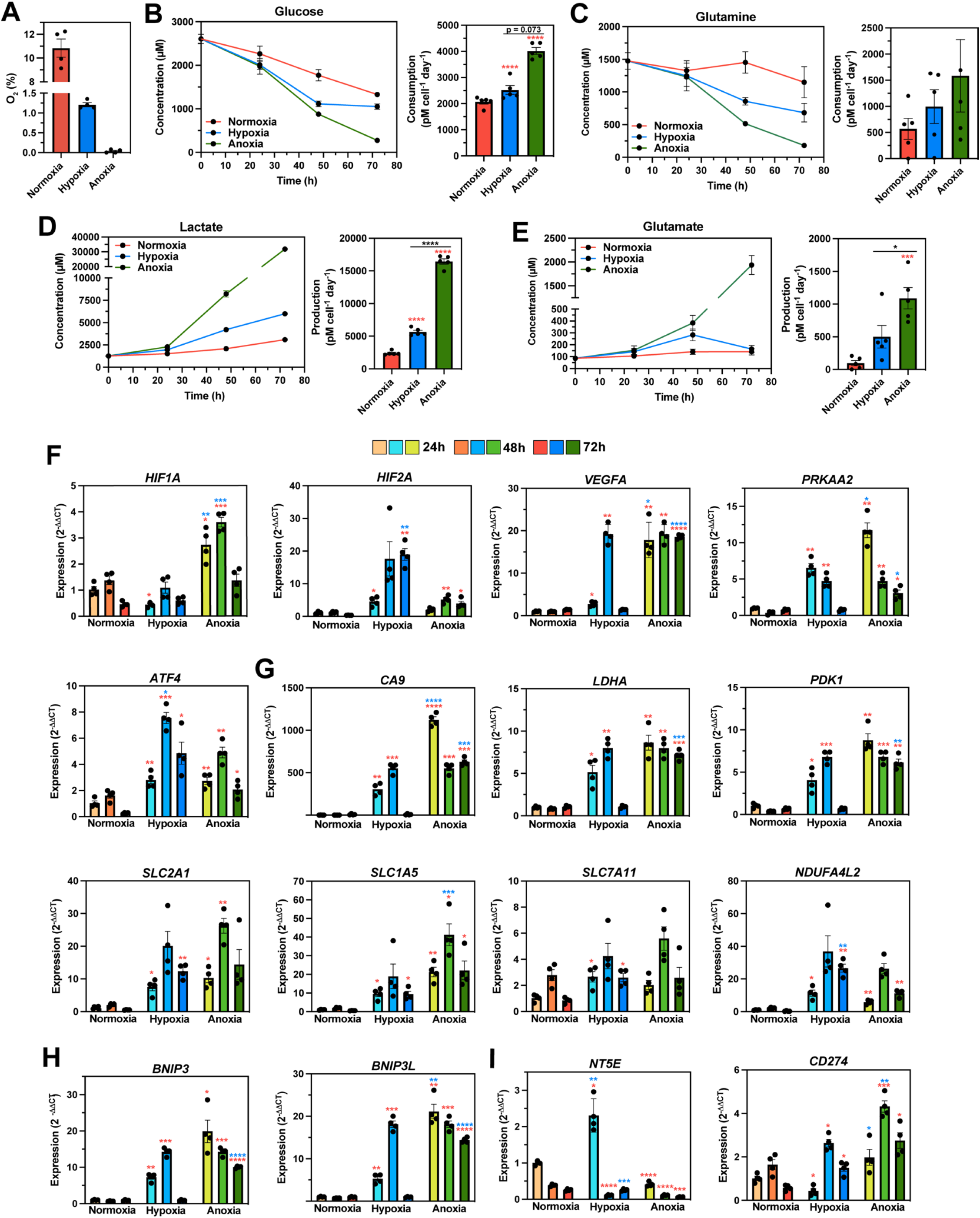
Pericellular anoxia induces stronger metabolic reprogramming and distinct transcriptional *HIFA* and HRE controlled gene responses compared to pericellular hypoxia in MCF7 cells. (A) Average pericellular O_2_ tensions in MCF7 cultures placed in 18.6%, 4.5%, and 1% O_2_ incubators for 72 hours. **(B–E)** Extracellular concentrations of glucose **(B)**, glutamine **(C)**, lactate **(D)**, and glutamate **(E)** in different O_2_ tensions for 72 hours. Normalized metabolite concentrations over time (left) and metabolite consumption or production rates for 72 hours (right). **(F–I)** Gene expression levels of genes associated with the hypoxic response **(F)**, metabolic reprogramming **(G)**, autophagy **(H)**, and immunosuppression **(I)** in different O_2_ tensions for 72 hours. Data were analyzed using ANOVA with Tukey’s correction. N = 4 biological samples per condition. Color code for asterix (*): Red colored * indicate comparison to control (normoxia) and blue colored * indicate comparison between hypoxia and anoxia.

### Pericellular hypoxia *vs.* anoxia induces distinct transcriptional HIFα and HRE gene responses in MCF7 cells

We examined MCF7 transcriptional responses to low O_2_ tensions at 24, 48, and 72 hours, including genes associated with (i) the hypoxic response (*e.g.*, hypoxia inducible factor 1 subunit alpha (*HIF1A*), hypoxia inducible factor 2 subunit alpha (*HIF2A*), vascular endothelial growth factor A (*VEGFA*), protein kinase AMP-activated catalytic subunit alpha 2 (*PRKAA2*), and activating transcription factor 4 (*ATF4*)); (ii) metabolic reprogramming (*e.g.*, carbonic anhydrase 9 (*CA9*), lactate dehydrogenase A (*LDHA*), pyruvate dehydrogenase kinase 1 (*PDK1*), solute carrier family 2 member 1 (*SLC2A1*, GLUT1), solute carrier family 1 member 5 (*SLC1A5*), solute carrier family 7 member 11 (*SLC7A11*, xCT), and NADH dehydrogenase 1 alpha subcomplex, 4-like 2 (*NDUFA4L2*)); (iii) mitophagy (*e.g.*, Bcl-2 interacting protein 3 (*BNIP3*) and Bcl-2 interacting protein 3 like (*BNIP3L*)); and (iv) the immunosuppressive tumor microenvironment (TME) (*e.g.*, cluster of differentiation 274 (*CD274*, PD-L1) and 5’-nucleotidase ecto (*NT5E*, CD73^49, 50^) (**Figure 5F–I**). Nearly all of these genes are direct HIF targets (*i.e.*, hypoxia-responsive element (HRE)-controlled genes), excluding *PRKAA2*, *ATF4*, and *SLC1A5*.^32, 42^

In anoxia, *VEGFA*, *CA9*, *PDK1*, *BNIP3*, and *BNIP3L* show elevated expression compared to normoxia throughout the time course. However, in hypoxia, these genes steadily increased expression and peaked after 48 hours, followed by a drop to normoxic expression levels after 72 hours. Interestingly, *HIFA* expression was different in hypoxia and anoxia: *HIF1A* had higher expression in anoxia (6- and 3-fold after 24 and 48 hours, respectively) and *HIF2A* had higher expression in hypoxia (5-fold after 72 hours) (**Figure 5F**). In addition, MCF7 had higher expression of *PRKAA2* (AMPK α2 subunit) in anoxia (2- and 4-fold after 24 and 72 hours, respectively), suggesting lower ATP availability in anoxia.^51^ Interestingly, hypoxia induced 2-fold higher expression of *ATF4* after 48 hours than did anoxia, indicating a stronger integrated stress response in hypoxia.^52^ Glycolytic genes had higher expression in anoxia (*LDHA* 7-fold after 72 hours and *PDK1* 9-fold after 72 hours). *SLC2A1* (GLUT1) expression levels also trended higher in anoxia throughout the time course. *SLC1A5* expression levels were 2-fold higher in anoxia after 48 hours. Overall, these results support the metabolic profiles in **Figure 5B–E**.

*BNIP3* and *BNIP3L* expression levels were higher in anoxia after 72 hours, suggesting an upregulation in mitophagy in anoxia.^42^ Lastly, in the context of immunosuppression, hypoxia induced 6-fold higher expression of *NT5E* (CD73, extracellular AMP to adenosine conversion^49, 50^) after 24 hours. Anoxia induces 2-fold higher expression of *CD274* (PD- L1) after 48 hours. Overall, these results indicate that hypoxia and anoxia induce distinct expression profiles in both *HIFA* and HRE responsive genes in MCF7.

### Proteomic characterization of the temporal differences between pericellular hypoxic and anoxic responses in 4T1 cells

After looking at transcriptional responses to hypoxia and anoxia, we aimed to better understand changes in protein expression in response to low O_2_ tensions. To this end, we applied plexDIA^53^ to understand how the proteome changes in response to hypoxia and anoxia in a murine triple negative breast cancer (TNBC) cell line (4T1). PCA of the proteome shows clustering by O_2_ tension and by day (**Figure S4A**). Furthermore, hypoxic and anoxic samples shift away from normoxic samples along PC1 over time, indicating continuing changes during low O_2_ responses.

For the hypoxic response, differential protein abundance analysis indicates no significantly upregulated or downregulated proteins after 1 day of culture, with the maximum response occurring after 3 days. On the other hand, the anoxic response had 50 upregulated and 5 downregulated proteins after day 1, and the response peaked after only 2 days. The number of changing proteins was higher in anoxia than hypoxia for all three days (**Figure S4B**).

Protein set enrichment analysis (PSEA) was performed to compare each low O_2_ response between days. In agreement with the differential protein abundance analysis, PSEA suggests that the anoxic response is faster and peaks by day 2: most of the changes occur for 2 days *vs.* 1 day (**Figure S4C**). For both O_2_ conditions, hypoxia-associated pathways are upregulated throughout the time course, including hypoxia, glycolysis, cholesterol homeostasis, fatty acid metabolism, and epithelial-to-mesenchymal transition (EMT). Interestingly, Myc targets were downregulated in both tensions for all days, which may contribute to the reduction in proliferation at low O_2_ tensions.^54^ Overall, this temporal characterization of the proteome suggests that the response to anoxia is stronger and faster than the response to hypoxia in 4T1 cells.

### Characterizing proteomic differences in metabolic reprogramming between pericellular hypoxic and anoxic responses in 4T1 cells

To further characterize low O_2_ responses in 4T1 cells, we explored protein synthesis, hypoxic responses, and metabolic reprogramming at the pathway and protein level. As expected, RNA processing and protein translation pathways were downregulated in hypoxia and anoxia compared to nomoxia (**Figure 6A**). Most of these pathways were upregulated in hypoxia compared to anoxia, suggesting downregulation as a function of O_2_ tension.Surprisingly, translation elongation was upregulated in anoxia compared to hypoxia after 1 day of culture, suggesting distinct regulation in the acute anoxic response.

**Figure 6:**
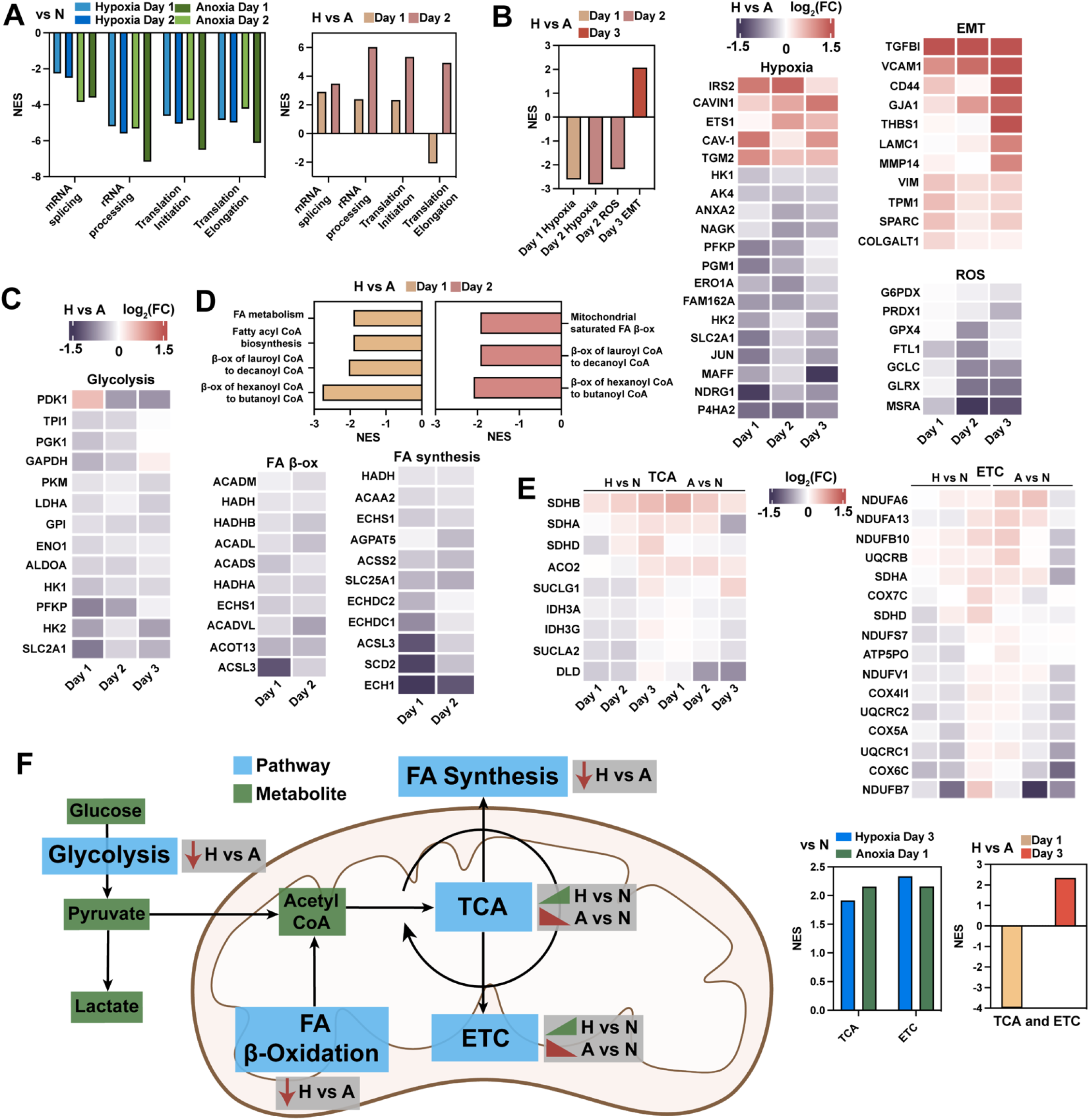
Proteomic characterization of pericellular hypoxic and anoxic metabolic reprogramming in 4T1 cells. (A) Protein set enrichment analysis (PSEA) using the Hallmark database of mRNA processing and protein translation pathways for Hypoxia (H) *vs.* Normoxia (N), Anoxia (A) *vs.* Normoxia, and Hypoxia *vs.* Anoxia. (B) PSEA using the Hallmark database of hypoxic response pathways (left). Heat maps of selected proteins in Hallmark hypoxia, epithelial to mesenchymal transition (EMT), and reactive O_2_ species (ROS) pathways (right) (Hypoxia *vs.* Anoxia). (C) Heat map of glycolytic and hypoxic response proteins. (D) PSEA using Reactome database for fatty acid (FA) metabolism pathways for Hypoxia *vs.* Anoxia (top). Heat maps of selected proteins in fatty acid β- oxidation and synthesis Reactome pathways (bottom). (E) Heat maps of selected proteins from Tricarboxylic acid (TCA) cycle and electron transport chain (ETC) processes for Hypoxia *vs.* Normoxia and Anoxia *vs.* Normoxia. PSEA using the Reactome database of TCA and ETC pathways for Hypoxia *vs.* Normoxia, Anoxia *vs.* Normoxia (left), and Hypoxia *vs.* Anoxia (right). NES = normalized enrichment score. N = 3 biological replicates and N = 2-3 technical replicates per condition.

Next, we looked at pathways associated with the hypoxic response, and discovered that hypoxia and reactive oxygen species (ROS) pathways were upregulated in anoxia and EMT was upregulated in hypoxia (**Figure 6B**). Proteins associated with hypoxia-induced stress responses showed higher abundance in anoxia for all three days. In addition, proteins associated with tumor progression, invasion, and metastasis, were upregulated in hypoxia for the duration of the experiment.

Figure 6C illustrates that 9 out of 10 enzymes involved in glycolysis are more abundant in anoxia compared to hypoxia. In addition, proteins involved in hypoxia-mediated metabolic reprogramming show increased abundance in anoxia. We also examined fatty acid β-oxidation and synthesis, which are downregulated and upregulated in hypoxia respectively.^32, 42^ Interestingly, PSEA suggests that both of these processes are downregulated in anoxia compared to hypoxia for the first 2 days of culture (Figure 6D). Most enzymes involved in mitochondrial acyl-CoA to acetyl-CoA conversion (*i.e.*, fatty acid β-oxidation) were upregulated in anoxia.

Finally, we examined oxidative phosphorylation, which is expected to decrease in low O_2_ tensions.^32, 42^ Unexpectedly, for hypoxia *vs.* anoxia, the tricarboxylic acid (TCA) cycle and electron transport chain (ETC) processes were downregulated after 1 day of culture and upregulated after 3 days of culture (Figure 6E). Clustering of TCA and ETC enzymes suggests increasing oxidative phosphorylation activity in hypoxia, yet decreasing activity in anoxia (compared to normoxia). Taken together, these results suggest that 4T1 cells have distinct hypoxic and anoxic metabolic responses (Figure 6F).

## Discussion

Despite the widespread use of O_2_-controlling chambers in hypoxia-related research, studies quantifying pericellular O_2_ concentrations and their impact on cellular response are surpisingly lacking. In the current study, we discovered vast differences between incubator setpoints and pericellular O_2_ tensions in every cell type tested. Our results highlight a major challenge with portable chambers and tri-gas incubators: cultures cannot be conditioned to start at the desired O_2_ tension due to rapid reoxygenation of media upon exposure to normoxia.

Without media conditioning, MCF7 cultures can take 1–14 hours to reach 1% O_2_ depending on experimental set-up. Not only does this time difference introduce significant variability between experiments, it suggests that shorter hypoxic experiments may not even reach hypoxia. We also show that changing cell culture parameters induce a sustained difference of HIF stabilization kinetics for at least 5 days for two different cell lines. Unless a hypoxic workstation with preconditioned media is used, pericellular O_2_ tension must be determined to report accurate O_2_-controlled incubation times.

Challenging the belief that incubators accurately control O_2_ for cell cultures, we discovered that pericellular anoxic tensions are common in both physioxic (5% O_2_) and hypoxic (1% O_2_) conditions due to cellular O_2_ consumption. This effect occurred in both primary human adherent and suspension cultures, with commonly used cell densities, medium volumes, and culture vessel types. Furthermore, our results suggest that physioxic cultures are routinely hypoxic. O_2_ tension in physioxia can vary greatly (0.1– 4.5% O_2_) depending on experimental set-up, engendering reproducibility concerns. These findings are a major concern for physioxic culture of stem cell expansion and differentiation^55–57^, and controlling pericellular O_2_ tension may improve our understanding of these processes.

Our pericellular O_2_ tension results for primary human hepatocyte cultures, a prominent model in drug metabolism, underpin how the incubator setpoint is a poor indicator of the O_2_ cells experience. Hepatocytes cultured in physioxia experienced anoxia for 24 hours, followed by a rapid reoxygenation upon media exchange. RNA-seq results indicate that O_2_ fluctuations in these conditions induce an upregulation in cellular responses to mitochondrial and NADPH oxidase ROS production (*e.g.*, superoxide, hydrogen peroxide).^58, 59^ Oxidative stress increased cell death via upregulation of TNF (apoptosis) and IL-1β (pyroptosis) production.^60, 61^ Damage-associated molecular patterns (DAMPs) released by dying cells activate toll-like receptor (TLR) and pattern recognition receptor (PRR) pathways, inducing key signatures of a sterile inflammatory response: complement activation, inflammasome complex assembly, MHC class II upregulation, and IL-6 production^62, 63^. During reoxygenation, hepatocytes increased mitochondria and ribosome biogenesis to meet ATP and protein translation demands as the cells recovered from hypoxic exposure.^42^ Ultimately, these results indicate that physioxic culture of hepatocytes drives a cellular response mimicking liver ischemia-reperfusion injury, a major risk factor in graft dysfunction in liver transplanation.^61^

We developed a reaction-diffusion model, which accurately predicted pericellular O_2_ tensions of MDA-MB-231 cultures at various incubator setpoints. This novel tool can design O_2_-controlled cell culture experiments, modulating parameters like the cell density, culture vessel type, or medium volume to achieve desired pericellular O_2_ concentrations. Our finding that the effect of cellular O_2_ consumption increases as culture vessel size decreases suggests that smaller vessels (*e.g.*, 96-well plates) should be avoided for O_2_- controlled cell culture. This observation carries significant implications for immune cell culture, which are typically done at high cell densities in small culture vessels. Future iterations of the model should consider incorporating cell growth rates as a function of O_2_ tension. In addition, experiments to determine O_2_ consumption rates (V_max_) for different cell types are needed. Such studies may uncover O_2_ consumption trends applicable to most cell types.

Using the reaction-diffusion model, we performed the first relation of pericellular O_2_ tension to biological response. We investigated the metabolic, transcriptomic, and translational responses to hypoxia (1–2% O_2_) and anoxia (0.0% O_2_) in two different breast cancer cell lines. Quantification of 14 genes associated with hypoxia (11 are direct HIF targets) and *HIF1A*/*HIF2A* suggest distinct transcriptional responses in hypoxia and anoxia. In anoxia, we found higher transcription in *HIF1A*, hypoxic markers (*VEGFA*, *CA9*), metabolism (*LDHA*, *PDK1*, *SLC1A5*), mitophagy (*BNIP3*, *BNIP3L*), and *CD274* (PD-L1) compared to hypoxia. Conversely, *HIF2A*, *ATF4*, *NDUFA4L2*, and *NT5E* (CD73) was upregulated in hypoxia. The increase in *HIF1A* transcription in anoxia may occur through ROS-induced PI3 kinase (PI3K) and protein kinase C (PKC) pathways^64, 65^, since our proteomic analysis suggests that ROS production is higher in anoxia.

A previous study reported that maximum HIF1 DNA-binding activity occurs at pericellular 0.5% O_2_ and sharply decreases as tension approaches 0% O_2_.^66^ This suggests that in anoxic culture, HIF is maximally stabilized as cultures approach anoxia (rather than in anoxia *per se*) inducing a strong HRE transcriptional response. Ultimately, 0.5% pericellular O_2_ may be the ideal setpoint for hypoxic cell culture, as long as tensions do not drop to anoxic levels. Future studies will examine the post-translational modification of both HIF1/HIF2 and downstream responses in different low O_2_ tensions (0 – 3% O_2_).

To further understand low O_2_ tension responses, we performed an in-depth characterization of the 4T1 proteome in response to low O_2_ tensions. This analysis suggests that the global translational response to anoxia is stronger and faster than the hypoxic response. Yet, the responses are distinct: we found an upregulation of EMT proteins in hypoxia and an increased ROS response in anoxia for 72h of culture. EMT is a critical step in hypoxia-driven metastasis^67^ and metastastic breast cancer represents the most advanced stage of the disease.^68^ Our findings suggest that mitochondrial dysfunction and ROS production are higher in anoxia than in hypoxia.^69^

Ultimately, our exploration of breast cancer responses to low O_2_ tensions suggest that anoxia is not suitable to model hypoxia. This is fortified by the fact that the median O_2_ tension in breast tumors is 10 mmHg (1.3% O_2_).^70^ Equally important, our findings uncover that breast cancer cells respond non-monotonically to low O_2_, since many aspects of the low O_2_ response are upregulated in hypoxia compared to anoxia. Future work will further explore these responses and relate them to hypoxic environments *in vivo*.

## Conclusion

O_2_ is a critical factor for mammalian bioenergetic homeostasis and serves as a substrate for over 200 enzymatic reactions.^12^ O_2_-controlled cell culture attempts to mimic O_2_ tensions that cells are exposed to *in vivo* and is therefore a critical tool for biological research. Herein, we report the discovery that the metric used to determine O_2_ concentration for *in vitro* cultures, the incubator setpoint, is a poor indicator of the O_2_ tension cells actually experience (*i.e.*, pericellular O_2_ tension) due to cellular O_2_ consumption. Standard physioxic (5% O_2_) and hypoxic (1% O_2_) protocols routinely induce anoxia (0.0% O_2_). Furthermore, in physioxic culture, pericellular O_2_ tension is highly dependent on cell culture parameters, making reproducibility difficult. Highlighting the significance of these findings, we demonstrated that a key drug metabolism model, primary human hepatocytes, undergo an effect similar to ischemia-reperfusion injury when cultured in physioxia. To address these challenges, we developed a reaction-diffusion model that predicts pericellular O_2_ tension *a priori*. Using this tool, we controlled pericellular O_2_ tension in two breast cancer models to explore transcriptional and translational responses to hypoxia and anoxia. We discovered that breast cancer cells respond non-monotonically to low O_2_. Overall, this work calls for a fundamental change to how O_2_-controlled cell culture is performed and suggests that *pericellular O_2_-controlled cell culture* is necessary to accurately model O_2_ tension.

## Methods

### *In vitro* O_2_ measurements

Adhesive optical O_2_ sensor spots (OXSP5-ADH-STER, PyroScience GmbH) were used to measure the O_2_ concentration of media and cell cultures. Sensors were placed on the culture vessel surface and a cable adapter (SPADBAS, PyroScience GmbH) was glued on the opposite side of the culture vessel (lined up with the sensor). Glue was allowed to dry overnight. Optical fiber cables (SPFIB-BARE, PyroScience GmbH) were placed within the adapters and connected to a computer via a meter (FireSting O_2_, PyroScience GmbH). The 100% O_2_ calibration was performed with aerated Dulbecco’s phosphate-buffered saline (DPBS), and the 0% O_2_ calibration was was performed using the factory calibration value. For cell culture experiments, cells were seeded in sensor-containing culture vessels and pericellular O_2_ was measured. A temperature probe (TDIP15, PyroScience GmbH) connected to the meter was placed inside the same incubator as the sensor-containing culture vessels. To measure the O_2_ concentration at the top of the media or cell culture wells, needle-like probes (OXROB10, PyroScience GmbH) were attached to a micromanipulator (MM33, PyroScience GmbH) and placed at the media– gas interface, such that the probes were submerged at the top layer of the media. Holes were drilled in plate lids to allow the probes to reach the media.

### Cell culture

Mycoplasma-free cell lines, MDA-MB-231 (ATCC HTB-26), MCF7 (ATCC HTB-22), 4T1 (ATCC CRL-2539), and primary mammary epithelial (ATCC PCS-600-010) were obtained from the American Type Culture Collection (ATCC). MDA-MB-231, MCF7, and 4T1 were maintained in Leibovitz’s L-15 medium (Cytiva), Eagles’ Minimum Essential Medium (EMEM) with L-glutamine (Quality Biological) and Dulbecco’s Minimum Essential Medium (DMEM) (Corning), respectively, with 10% fetal bovine serum (FBS, Corning) and 1% penicillin/streptomycin (P/S, Invitrogen). Mammary epithelial cells were cultured in basal medium (ATC PCS-600-030) with cell growth kit (ATCC PCS-600-040). MCF7 and MDA-MB-231 hypoxia-inducible factor (HIF) reporter cell lines were transduced and selected as previously described.^28^ For all cultures, passage number did not exceed 20.

Human dendritic cells (DCs) were differentiated from cryopreserved CD14+ monocytes over a period of 7 days as previously described.^71^ In brief, monocytes were seeded into a 6-well plate in ImmunoCult™-ACF Dendritic Cell Medium (Stemcell Technologies), supplemented with recombinant human granulocyte-macrophage colony-stimulating factor (GM-CSF) (50 ng/mL) (R&D Systems) and recombinant human interleukin-4 (IL-4) (50 ng/mL) (R&D systems).

Primary human hepatocytes (HUCPG, Lonza) were cultured following the manufacturer’s protocol. Briefly, hepatocytes were thawed in thawing medium (MCHT50, Lonza), seeded onto a 24-well plate (BioCoat Collagen I, Corning) with plating medium (MP100 and MP250, Lonza), then cultured with maintenance medium (CC-3198, Lonza), which was exchanged after 24 hours.

For O_2_-controlled experiments, cells were incubated within a tri-gas incubator (Heracell VIOS 160i, Thermo Fisher Scientific) which was kept closed during the duration of the experiment.

### Cellular hypoxia kinetics

MCF7 cells were seeded onto 60 mm glass dishes (Cellvis) overnight and incubated with Hoechst 33342 (NucBlue Live ReadyProbes Reagent, Invitrogen), 10 µM Image-iT^TM^ Red Hypoxia Reagent (Invitrogen), and 1 µM Celltracker Orange for 30 minutes at 37°C. Cells were placed within a humidified incubator at 37°C, 5% CO_2_, and 1% O_2_/99% N2 (O_2_ Module S, CO_2_ Module S, and Temp Module S, Zeiss) attached to a confocal microscope (LSM 880 with Airyscan, Zeiss). Images were taken every 30 minutes for 12 hours. Average fluorescence per cell was calculated using Fiji. Briefly, cell area multiplied by background flourescence was subtracted from the cell’s integrated density. At least thirty cells were analyzed per image.

### Hypoxia-inducible factor (HIF) stabilization kinetics

For cell density experiments, MCF7 reporter cells were seeded into a 12-well plate at different densities (7,000 and 29,000 cells/cm^2^) overnight. Cells were cultured at 1% O_2_ for 4 days, incubated with Hoechst 33342 and confocal images were taken daily. Percentage of GFP+ among MCF7 reporter cells was determined using Fiji (Threshold and Analyze Particles). For culture vessel type experiments, MDA-MB-231 reporter cells were seeded onto a 24-well plate or T25 flask (30,000 cells/cm^2^) and cultured at 1% O_2_ for 5 days. Flow cytometry (Cytoflex S, Beckman Coulter) of live singlets was used to determine the GFP positive fraction.

### Mitochondrial inhibition

MCF7 were seeded overnight in a 24-well plate and incubated at 5% O_2_ for 24h. 10 µL of DPBS or sodium azide (Sigma Aldrich) (final concentration = 5 mM) were added into cultures. For cytotoxicity studies, MCF7 were seeded overnight and incubated with ± 5 mM sodium azide for 6 hours at 5% O_2_. Cells were incubated with 1:1000 live/dead dye (LIVE/DEAD Fixable Kit, Thermo Fisher Scientific) for 1 hour. Flow cytometry (Attune NxT, Thermo Fisher Scientific) of live singlets was used to determine the cell viability.

### Cell viability time course

MCF7 were seeded into a T25 flask (20,000 cells/cm^2^) overnight and cultured at 18.6%, 5%, or 1% O_2_ for 6 days. Every 2 days, media was harvested, and trypan blue staining and a hemocytometer were used to determine live and dead detached cells. Attached cells were trypsinized (Trypsin-EDTA, Gibco) and counted using the same method.

### Library preparation with polyA selection and Illumina sequencing

RNA was extracted from hepatocytes (NucleoSpin RNA, Macherey-Nagel) and quantified using a Qubit 2.0 Fluorometer (Life Technologies). RNA integrity was checked using Agilent TapeStation 4200 (Agilent Technologies). RNA sequencing libraries were prepared using the NEBNext Ultra II RNA Library Prep for Illumina per the manufacturer’s protocol (New England Biolabs). Briefly, mRNAs were enriched with Oligod(T) beads. Enriched mRNAs were fragmented for 15 minutes at 94 °C. First strand and second strand cDNA were subsequently synthesized. cDNA fragments were end repaired and adenylated at 3’ ends, and universal adapters were ligated to cDNA fragments, followed by index addition and library enrichment by PCR with limited cycles. The sequencing libraries were validated on the Agilent TapeStation 4200 and quantified by using Qubit 2.0 Fluorometer as well as by quantitative PCR (KAPA Biosystems). The sequencing libraries were multiplexed and clustered onto a flowcell. After clustering, the flowcell was loaded onto the Illumina instrument (HiSeq 400 or equivalent) according to the manufacturer’s instructions. The samples were sequenced using a 2x150bp Paired End (PE) configuration. Image analysis and base calling were conducted by the HiSeq Control Software (HCS). Raw sequence data (.bcl files) generated from Illumina HiSeq were converted into FASTQ files and de-multiplexed using Illumina bcl2fastq 2.20 software. One mismatch was allowed for index sequence identification. FASTQ files were trimmed with Trimmomatic^72^ and the quality was analyzed with FastQC.^73^ The human genome (GRCh38.p14) was annotated and reads were aligned using STAR.^74^ Gene counts were determined using FeatureCounts.^75^ Differential gene expression analysis was performed using DESeq2.^76^ Gene set enrichment analysis (GSEA)^77^ was performed using the clusterProfiler package in R.^78^

### Reaction-diffusion model

The unsteady state diffusion equation (1) with initial and boundary conditions (2–4) were used to describe O_2_ transfer between cell culture media and gas phase, where C is the concentration of O_2_, D is the diffusivity coefficient, and kLa is the mass transfer coefficient. x = 0 is the bottom of the well and x = L is the media height. A diffusivity coefficient of 0.09684 cm^2^/hr was used^79^ and experimental diffusion data were used to determine kL values. An analytical solution was determined (5–6). Michaelis–Menten kinetics were used for the reaction-diffusion model^38^ (7), where V_max_ is the maximum O_2_ consumption rate and Km is the O_2_ concentration at which the reaction rate is half of V_max_. Numerical values were determined using the MATLAB PDE solver (MathWorks).

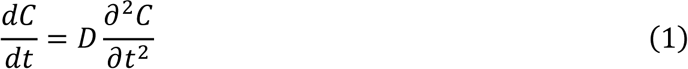

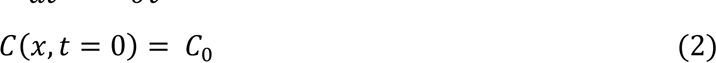

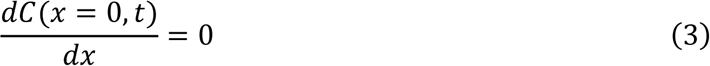

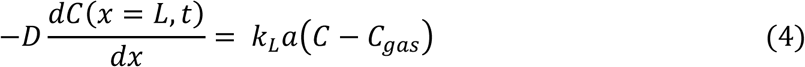

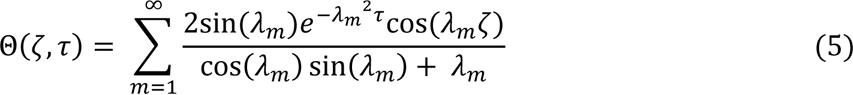

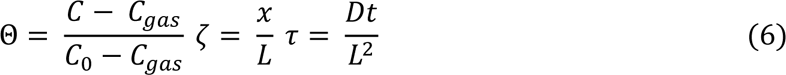

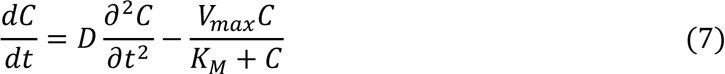

### Extracellular metabolite quantification

Media was removed from cell cultures and centrifuged at 250g for 5 minutes. Resulting supernatant was stored at -20⁰C and used for metabolite quantification. Live cells from cultures were determined using trypan blue staining and a hemocytometer. Glucose uptake and lactate secretion were quantified by an Agilent 1260 high performance liquid chromatography (HPLC) Infinity II System equipped with a BioRad Aminex HPX-87H ion exchange column (300mm × 7.8 mm) operated at 60⁰C with a refractive index detector (RID) operated at 50⁰C.^80, 81^ The mobile phase was 14mM sulfuric acid with a flow rate of 0.6 mL/min. The injection volume of each sample was 10 μL. Peak areas for each compound for concentrations ranging from 0.125 g/L to 5 g/L were used to make calibration curves in OpenLab ChemStation (LTS 01.11) and then used to calculate metabolite quantifications. Glutamate and glutamine concentration was determined using the Glutamine/Glutamate-Glo Assay (Promega). Metabolite concentration was normalized by the cell number at each time point.

### Reverse transcription and quantitative PCR (RT-qPCR)

RT-qPCR was performed as previously described.^82, 83^ RNA was extracted from MCF7 cultures (NucleoSpin RNA, Macherey-Nagel) and the quality was checked using a NanoDrop One spectrophotometer (Thermo Fisher Scientific). Reverse transcription was conducted using a High-Capacity cDNA Reverse Transcription Kit (Applied Biosystems) on a MyCycler thermal cycler (Bio-Rad). Gene expression was quantified using the following TaqMan Gene Expression Assays (Thermo Fisher Scientific) on an MX3005P QPCR System (Agilent Technologies): *VEGF-A* (Hs00900055_m1), *CA9* (Hs00154208_m1), *LDHA* (Hs01378790_g1), *PDK1* (Hs01561847_m1), *NT5E* (Hs00159686_m1), *PRKAA2* (Hs00178903_m1), *BNIP3L* (Hs00188949_m1), *BNIP3* (Hs00969291_m1), *HIF1A* (Hs00153153_m1), *HIF2A* (Hs00909569_g1), *SLC2A1* (Hs00892681_m1), *SLC1A5* (Hs01056542_m1), *SLC7A11* (Hs00921938_m1), *NDUFA4L2* (Hs00220041_m1), *BNIP3* (Hs00969291_m1), *BNIP3L* (Hs00188949_m1), *NT5E* (Hs00159686_m1), *CD274* (Hs00204257_m1), and *ACTB* (Hs01060665_g1).

### Bulk proteomic sample preparation

Harvested cells were washed with PBS and resuspended in Mass Spectrometry grade Water (Fisher Scientific, W6500) and frozen at -80°C. Cells were lysed by heating at 90°C for 10 mins^84^, protein concentrations for each lysate were measured using a Nanodrop (A205). Proteins were digested to peptides per the SCoPE2 protocol.^85, 86^ Briefly, 10 µg of protein per sample were digested in a solution containing 100mM triethylammonium bicarbonate at pH 8.5 (TEAB) (Sigma Aldrich, T7408), benzonase nuclease (Millipore Sigma, Cat E1014) and Trypsin Gold (Promega, V5280). The protease was added at a 1:20 enzyme to substrate ratio and LC-MS grade water was added to maintain its concentration at 20 ng/uL. The reaction was carried out for 12 hours at 37°C. Digested peptides were subsequently dried down in a SpeedVac vacuum evaporator and resuspended in 200mM TEAB (pH 8.5). Samples were randomized and labelled using either d0, d4 or d8 of mTRAQ mass tags (SciEx, 4440015, 4427698, and 4427700) in a reaction that maintained 1/3rd organic phase and the manufacturer’s suggested reagent to peptide ratio (1U for 100 µg of peptides). The labelling was carried out for 2 hours at room temperature and excess, unreacted label was quenched by adding hydroxylamine (Sigma Aldrich, 467804) to 0.2% v/v and leaving at room temperature for 1 hour. Two samples from each label were randomly selected and 50 nng analyzed in data-dependent acquisition (DDA) mode to evaluate labelling efficiency.

Samples from each label were combined in equal amounts to make a plexDIA^53^ set that was dried down and resuspended in 0.1% formic acid (Thermo Fisher, 85178) in MS grade water to a final concentration of 1 µg/µL. Samples within a plexDIA set were randomly paired; a few samples across labels were repeated across multiple sets.

### Proteomics data acquisition

The separation was performed at a constant flow rate of 200 nL/min using a Dionex UltiMate 3000 UHPLC, and 1 µL of sample was loaded onto a 25cm x 75uM IonOpticks Odyssey Series column (ODY3-25075C18). The separation gradient was 4% buffer B (80% acetonitrile in 0.1% Formic Acid) for 11.5 minutes, a 30 second ramp up to 12%B followed by a 63 minute linear gradient up to 32%B. Subsequently, buffer B was ramped up to 95% over 2 minutes and maintained as such for 3 additional minutes. Finally, buffer B was dropped to 4% in 0.1 minutes and maintained for 19.9 additional minutes.

The mass spectra were analyzed using a Thermo Scientific Q-Exactive mass spectrometer from minutes 20 to 95 of the LC method. An electrospray voltage of 1700V was applied at the liquid-liquid junction of the analytical column and transfer line. The temperature of the ion transfer tube was 250°C, and the S-lens RF level was set to 30.

Bulk data was collected in Data Independent Acquisition (DIA) mode, the duty cycle consisted of a total of 3 MS1 scans and 30 MS2 scans. All MS1 scans were conducted at 140,000 resolving power with a maximum injection time of 300 milliseconds and a target AGC of 3e6 with a scan range covering 378-1290 m/z. All MS2 scans were conducted at 35,000 resolving power, a maximum inject time of 110 ms, AGC target of 3e6 and normalized collision energy of 27. MS2 scans had variable isolation widths: 10 MS2 scans of 17 m/z isolation width (isolation window) followed the first and second MS1 scan respectively, the third MS1 was followed by 5 windows of 33 m/z, 2 windows of 40 m/z, 1 window of 80 m/z and a final window of 120 m/z.

### Proteomics data processing

DIA-NN^87^ (version 1.8.1) was used to search the raw files from each run. A predicted spectral library was made using the swissprot mouse FASTA database and in silico labelled to have mTRAQ as a fixed mod (+140.0949630177) on each trypsin digested peptide.

Peak height was used for quantification with a scan window of 1, mass accuracy of 10 ppm and MS1 accuracy of 5 ppm, MBR was enabled and search outputs were filtered at 1% Q value. The following commands were employed by use of the additional commands dialogue: –fixed-mod mTRAQ 140.0949630177, nK, –channels mTRAQ, 0, nK, 0:0; mTRAQ, 4, nK, 4.0070994:4.0070994; mTRAQ, 8, nK, 8.0141988132:8.0141988132*}*, –peak-translation, –ms1-isotope-quant, -ms1-base-profile, -ms1-subtract 2.

The report file containing filtered peptide level output was processed using R. First, the peptide level data was collapsed/summarized to a run x protein matrix using the diannmaxlfq function from the ‘diann’ R package.^87^ Subsequently, the matrix was log2 transformed and the protein levels in each run were normalized for differential loading amounts by adding to each protein value, the median of the difference between the vector of protein levels for that run to the vector of median values across all runs. Relative protein levels were obtained by subtracting away the mean value across runs for each protein.

In order to correct for biases specific to each mTRAQ label, kNN imputation (k = 3) was performed and ComBat^88^ was used with mTRAQ labels as batch covariates. Post batch correction two matrices were used for further analysis, one with imputed values and the other where the imputed values had been set back to NA. Differential protein expression analysis was performed using limma.^89^ Protein set enrichment analysis (PSEA)^77^ was performed using the clusterProfiler^78^ package in R.

### Statistical analysis

All data were presented as the mean ± standard error of the mean (SEM). Statistical analyses were performed using Prism 9 software (GraphPad). The statistical tests used are outlined in the figure captions. Values represent the mean ± standard error of the mean. Significance levels are reported as *p < 0.05, **p < 0.01, ***p < 0.001, ****p < 0.0001.

## Acknowledgments

The authors thank the Institute for Chemical Imaging and Living Systems (CILS) for their technical assistance with the confocal microscope and flow cytometer. The authors would like to thank Dr. James Sherley for his insightful feedback throughout this project. S.A.B gratefully acknowledges the financial support from the National Institutes of Health (1R01EB027705) and the National Science Foundation (DMR-1847843 and TI-2141019).

## Author Contributions

Z.J.R., T.C., and S.A.B. conceived the presented idea and designed the experiments. Z.J.R. conducted the experiments unless otherwise stated. T.C. conducted the MCF7 transcription analysis. S.K. conducted the proteomics experiments and N.S. supported the analysis. K.B. and A.N. conducted the dendritic cell experiments. B.W. supported the development of the computational model. G.Z. conducted the glucose and lactate quantitation. C.T. and D.G. helped design the kinetic and pericellular measurement experiments and supported the transcriptional and translational analysis. Z.J.R. wrote the manuscript and generated the figures. All authors discussed the results, commented on, and proofread the manuscript. The principal investigator is S.A.B.

## Supplementary Information

**Figure S1:**
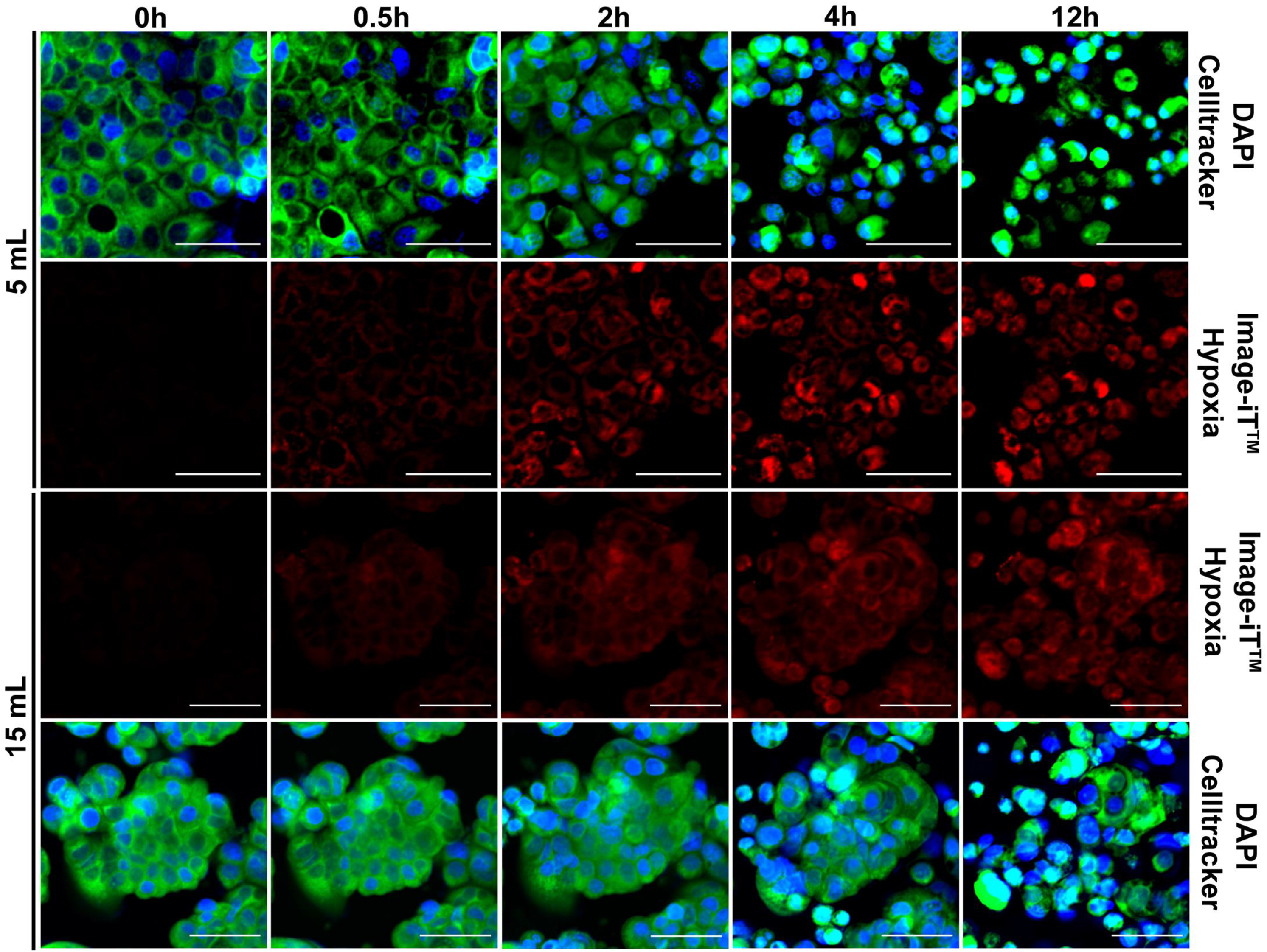
Medium volume influences cellular hypoxia kinetics in 1% O_2_ culture. Representative confocal images of MCF7 cultures in 60mm dishes with 5 mL or 15 mL of medium placed inside a 1% O_2_ chamber for 12h. Confocal images were taken every 30 minutes. Red = Image-iT^TM^ Hypoxia as an indicator of cellular hypoxia. Blue = nuclei stained with DAPI. Green = cytoplasm stained with Celltracker. N = 3 biological replicates per condition.

**Figure S2:**
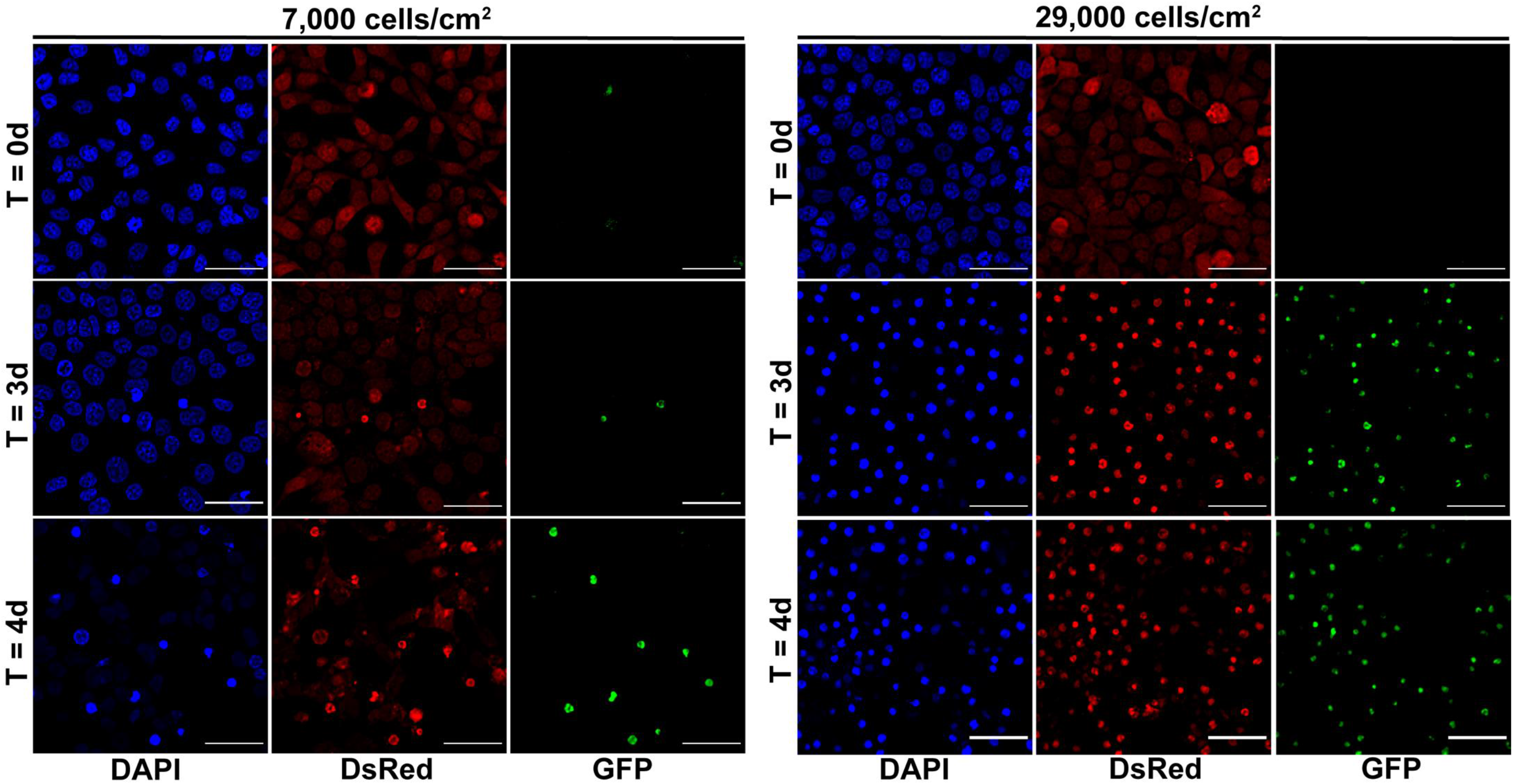
Cell density influences HIF stabilization kinetics in 1% O_2_ culture. Representative confocal images of MCF7 HIF reporter cells cultured at different densities (7,000 and 29,000 cells/cm^2^) in a 1% O_2_ incubator for 4 days. Images were taken at 0, 3 and 4 days. Different replicates were used for each time point to prevent reoxygenation. Blue = nuclei stained with DAPI. Red = Dsred (HIF-). Green = GFP (HIF+). N = 4 biological replicates per condition.

**Figure S3:**
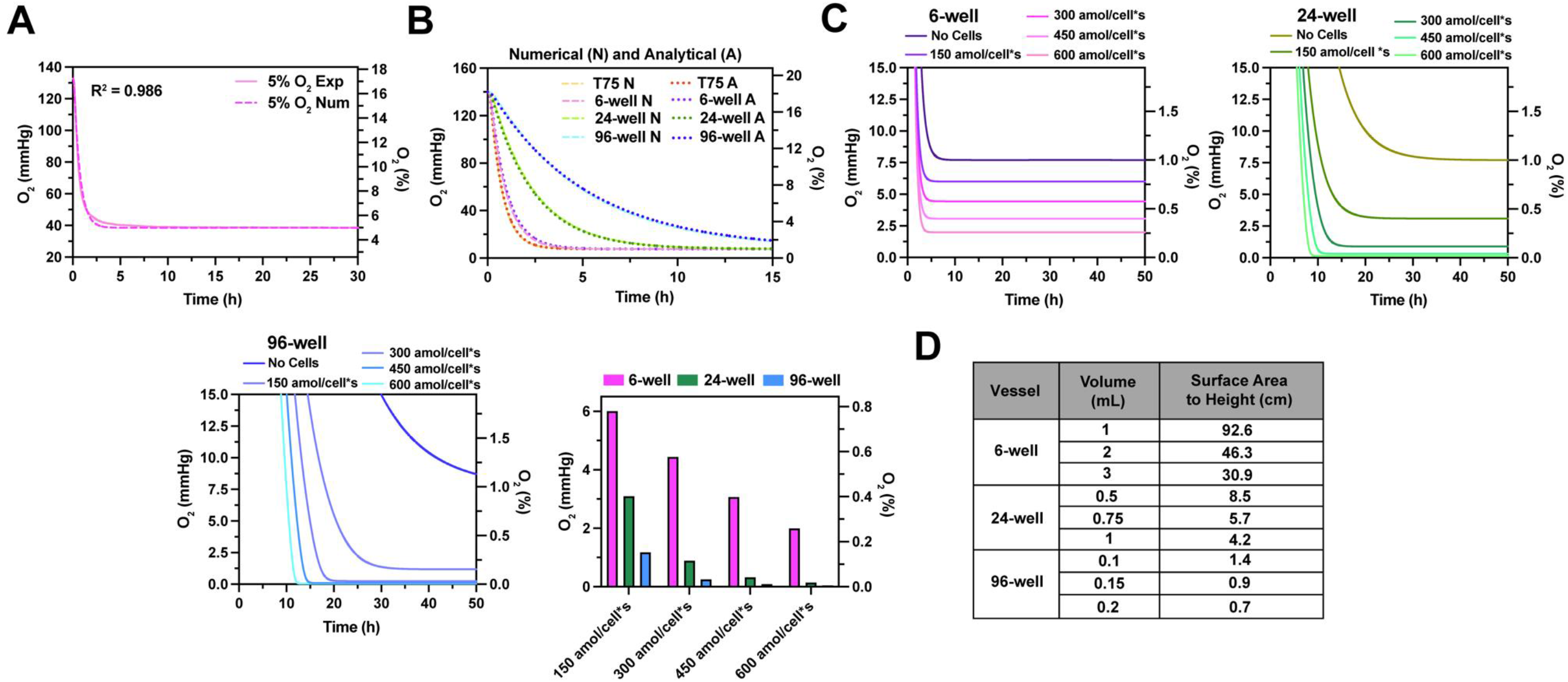
Developing a reaction-diffusion model to predict pericellular O_2_ tension in cell culture. (A) Experimental (solid) and numerical (dashed) O_2_ kinetics of media (EMEM + 10% FBS + 1% P/S) placed in a 5% O_2_ incubator. (B) Numerical (N) (dashed) and analytical (A) (dotted) O_2_ kinetics for media in different culture vessel types placed in a 1% O_2_ incubator. (C) Reaction-diffusion model predictions of cells (20,000 cells/cm^2^) with different O_2_ consumption rates (V_max_) in a 6-well, 24-well, or 96-well plate in a 1% O_2_ incubator. (D) Surface area to height ratios for 6-well, 24-well, or 96-well plates given commonly used volumes for each culture vessel.

**Figure S4:**
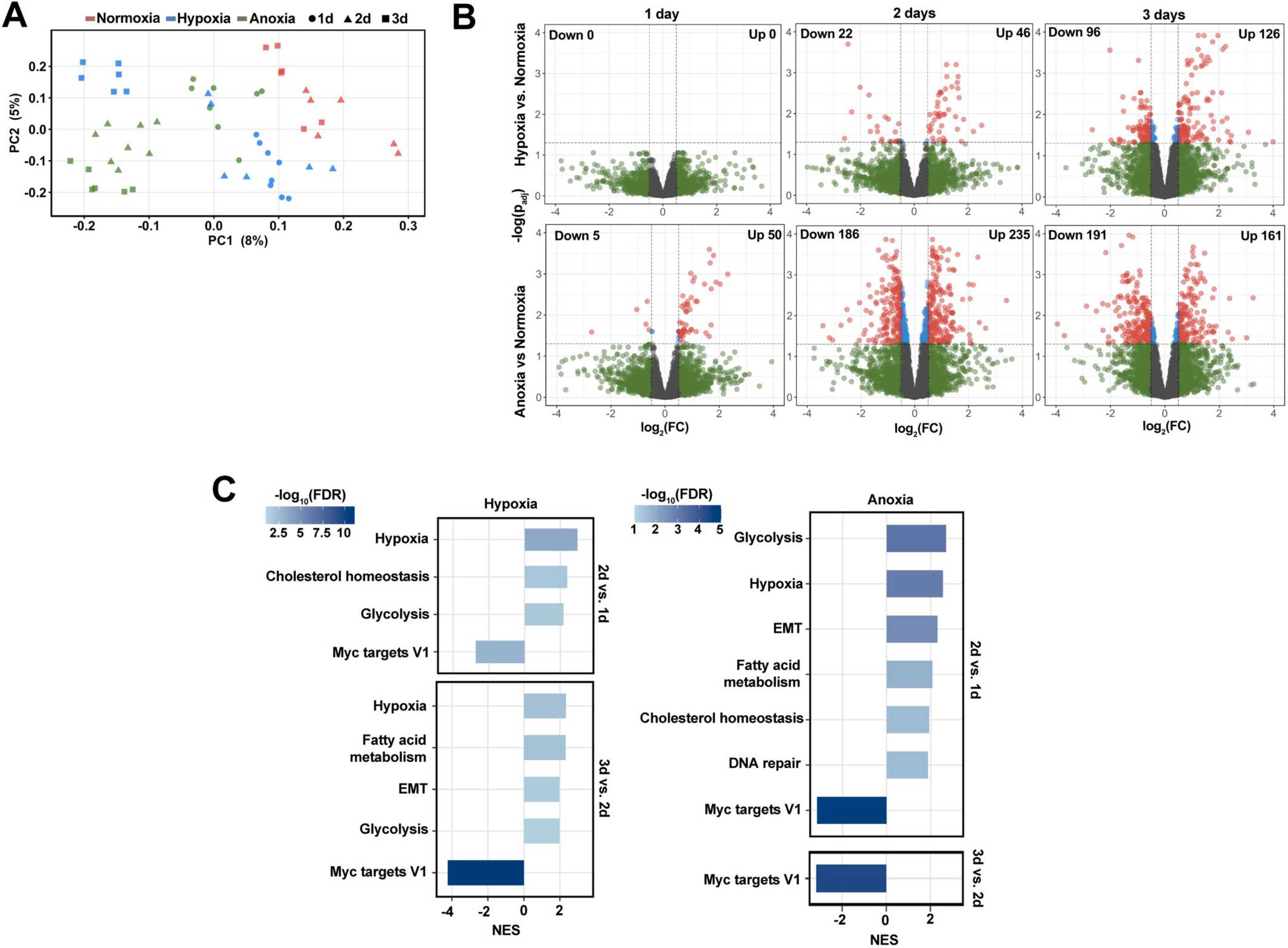
Proteomic characterization of the temporal differences between pericellular hypoxic and anoxic responses in 4T1. (A) Principal component analysis (PCA) of protein abundances for 4T1 cells cultured in different O_2_ tensions for 1, 2 or 3 days of culture. **(B)** Volcano plots indicating signficant downregulated or upregulated proteins (padj < 0.05 and |log2FC| ≥ 0.5) for Hypoxia *vs*. Normoxia (top) or Anoxia *vs*. Normoxia (bottom) for 1 day, 2 days, or 3 days of culture. **(C)** PSEA using the Hallmark database, comparing different days in hypoxia (left) or anoxia (right). N = 6 – 8 replicates per condition. NES = normalized enrichment score. N = 3 biological replicates and N = 2- 3 technical replicates per condition.

## References

1. Breslin S, O’Driscoll L. Three-dimensional cell culture: the missing link in drug discovery. Drug Discovery Today. 2013/03/01/ 2013;18(5):240–249. 10.1016/j.drudis.2012.10.003

2. Langhans SA. Three-Dimensional in Vitro Cell Culture Models in Drug Discovery and Drug Repositioning. 10.3389/fphar.2018.00006. Frontiers in Pharmacology. 2018;9:6.

3. Wang X, Rivière I. Manufacture of tumor- and virus-specific T lymphocytes for adoptive cell therapies. Cancer Gene Therapy. 2015/02/01 2015;22(2):85–94. doi:10.1038/cgt.2014.81

4. Hunsberger JG, Shupe T, Atala A. An Industry-Driven Roadmap for Manufacturing in Regenerative Medicine. Stem Cells Transl Med. 2018;7(8):564–568. doi:10.1002/sctm.18-0060

5. Lou J, Mooney DJ. Chemical strategies to engineer hydrogels for cell culture. Nature Reviews Chemistry. 2022/08/30 2022;doi:10.1038/s41570-022-00420-7

6. Colombani T, Rogers ZJ, Bhatt K, et al. Hypoxia-inducing cryogels uncover key cancer-immune cell interactions in an oxygen-deficient tumor microenvironment. Bioactive Materials. 2023/11/01/ 2023;29:279-295. 10.1016/j.bioactmat.2023.06.021

7. Lou J, Mooney DJ. Chemical strategies to engineer hydrogels for cell culture. Nature Reviews Chemistry. 2022/10/01 2022;6(10):726-744. doi:10.1038/s41570-022-00420-7

8. Pașca SP. The rise of three-dimensional human brain cultures. Nature. 2018/01/01 2018;553(7689):437-445. doi:10.1038/nature25032

9. LeSavage BL, Suhar RA, Broguiere N, Lutolf MP, Heilshorn SC. Next-generation cancer organoids. Nat Mater. Feb 2022;21(2):143–159. doi:10.1038/s41563-021-01057-5

10. Grönholm M, Feodoroff M, Antignani G, Martins B, Hamdan F, Cerullo V. Patient-Derived Organoids for Precision Cancer Immunotherapy. Cancer Res. Jun 15 2021;81(12):3149–3155. doi:10.1158/0008-5472.Can-20-4026

11. Ast T, Mootha VK. Oxygen and mammalian cell culture: are we repeating the experiment of Dr. Ox? Nature Metabolism. 2019/09/01 2019;1(9):858-860. doi:10.1038/s42255-019-0105-0

12. Flanigan WR, Jain IH. The Goldilocks Oxygen Principle: not too little and not too much. Nature Cardiovascular Research. 2022/12/01 2022;1(12):1101-1103. doi:10.1038/s44161-022-00178-7

13. Alva R, Mirza M, Baiton A, et al. Oxygen toxicity: cellular mechanisms in normobaric hyperoxia. Cell Biology and Toxicology. 2023/02/01 2023;39(1):111-143. doi:10.1007/s10565-022-09773-7

14. Sies H, Berndt C, Jones DP. Oxidative Stress. Annual Review of Biochemistry. 2017/06/20 2017;86(1):715-748. doi:10.1146/annurev-biochem-061516-045037

15. Timpano S, Guild BD, Specker EJ, et al. Physioxic human cell culture improves viability, metabolism, and mitochondrial morphology while reducing DNA damage. Faseb j. Apr 2019;33(4):5716–5728. doi:10.1096/fj.201802279R

16. Maddalena LA, Selim SM, Fonseca J, Messner H, McGowan S, Stuart JA. Hydrogen peroxide production is affected by oxygen levels in mammalian cell culture. Biochem Biophys Res Commun. Nov 4 2017;493(1):246–251. doi:10.1016/j.bbrc.2017.09.037

17. Baik AH, Haribowo AG, Chen X, et al. Oxygen toxicity causes cyclic damage by destabilizing specific Fe-S cluster-containing protein complexes. Mol Cell. Mar 16 2023;83(6):942–960.e9. doi:10.1016/j.molcel.2023.02.013

18. Ast T, Mootha VK. Oxygen and mammalian cell culture: are we repeating the experiment of Dr. Ox? Nat Metab. Sep 2019;1(9):858–860. doi:10.1038/s42255-019-0105-0

19. Atkuri KR, Herzenberg LA, Niemi A-K, Cowan T, Herzenberg LA. Importance of culturing primary lymphocytes at physiological oxygen levels. Proceedings of the National Academy of Sciences. 2007;104(11):4547. doi:10.1073/pnas.0611732104

20. Parrinello S, Samper E, Krtolica A, Goldstein J, Melov S, Campisi J. Oxygen sensitivity severely limits the replicative lifespan of murine fibroblasts. Nature Cell Biology. 2003/08/01 2003;5(8):741-747. doi:10.1038/ncb1024

21. Busuttil RA, Rubio M, Dollé ME, Campisi J, Vijg J. Oxygen accelerates the accumulation of mutations during the senescence and immortalization of murine cells in culture. Aging Cell. Dec 2003;2(6):287–94. doi:10.1046/j.1474-9728.2003.00066.x

22. Murphy CL, Polak JM. Control of human articular chondrocyte differentiation by reduced oxygen tension. 10.1002/jcp.10481. Journal of Cellular Physiology. 2004/06/01 2004;199(3):451–459.

23. Yazdani M. Technical aspects of oxygen level regulation in primary cell cultures: A review. Interdiscip Toxicol. Dec 2016;9(3-4):85–89. doi:10.1515/intox-2016-0011

24. Keeley TP, Mann GE. Defining Physiological Normoxia for Improved Translation of Cell Physiology to Animal Models and Humans. Physiol Rev. Jan 1 2019;99(1):161–234. doi:10.1152/physrev.00041.2017

25. Strey CW, Gestrich J, Beckhaus T, et al. Hypoxia and reoxygenation of primary human hepatocytes induce proteome changes of glucose metabolism, oxidative protection and peroxisomal function. Int J Mol Med. 2010/10/01 2010;26(4):577-584. doi:10.3892/ijmm_00000502

26. Joe NS, Wang Y, Oza HH, et al. Mebendazole Treatment Disrupts the Transcriptional Activity of Hypoxia-Inducible Factors 1 and 2 in Breast Cancer Cells. Cancers (Basel). Feb 20 2023;15(4)doi:10.3390/cancers15041330

27. Xie H, Song J, Godfrey J, et al. Glycogen metabolism is dispensable for tumour progression in clear cell renal cell carcinoma. Nature Metabolism. 2021/03/01 2021;3(3):327-336. doi:10.1038/s42255-021-00367-x

28. Godet I, Shin YJ, Ju JA, Ye IC, Wang G, Gilkes DM. Fate-mapping post-hypoxic tumor cells reveals a ROS-resistant phenotype that promotes metastasis. Nature Communications. 2019/10/24 2019;10(1):4862. doi:10.1038/s41467-019-12412-1

29. Jain IH, Calvo SE, Markhard AL, et al. Genetic Screen for Cell Fitness in High or Low Oxygen Highlights Mitochondrial and Lipid Metabolism. Cell. 2020/04/30/ 2020;181(3):716- 727.e11. 10.1016/j.cell.2020.03.029

30. Joe NS, Wang Y, Oza HH, et al. Mebendazole Treatment Disrupts the Transcriptional Activity of Hypoxia-Inducible Factors 1 and 2 in Breast Cancer Cells. Cancers. 2023;15(4). doi:10.3390/cancers15041330

31. Taylor CT, Colgan SP. Regulation of immunity and inflammation by hypoxia in immunological niches. Nature Reviews Immunology. 2017/12/01 2017;17(12):774-785. doi:10.1038/nri.2017.103

32. Taylor CT, Scholz CC. The effect of HIF on metabolism and immunity. Nature Reviews Nephrology. 2022/09/01 2022;18(9):573-587. doi:10.1038/s41581-022-00587-8

33. Fandrey J, Schödel J, Eckardt K-U, Katschinski DM, Wenger RH. Now a Nobel gas: oxygen. Pflügers Archiv - European Journal of Physiology. 2019/12/01 2019;471(11):1343-1358. doi:10.1007/s00424-019-02334-8

34. Wicks EE, Semenza GL. Hypoxia-inducible factors: cancer progression and clinical translation. The Journal of Clinical Investigation. 06/01/ 2022;132(11)doi:10.1172/JCI159839

35. McLimans WF, Crouse EJ, Tunnah KV, Moore GE. Kinetics of gas diffusion in mammalian cell culture systems. I. Experimental. Biotechnology and Bioengineering. 1968/11/01 1968;10(6):725-740. 10.1002/bit.260100603

36. McLimans WF, Blumenson LE, Tunnah KV. Kinetics of gas diffusion in mammalian cell culture systems. II. Theory. Biotechnology and Bioengineering. 1968/11/01 1968;10(6):741-763. 10.1002/bit.260100604

37. Peniche Silva CJ, Liebsch G, Meier RJ, Gutbrod MS, Balmayor ER, van Griensven M. A New Non-invasive Technique for Measuring 3D-Oxygen Gradients in Wells During Mammalian Cell Culture. Front Bioeng Biotechnol. 2020;8:595–595. doi:10.3389/fbioe.2020.00595

38. Wagner BA, Venkataraman S, Buettner GR. The rate of oxygen utilization by cells. Free Radic Biol Med. Aug 1 2011;51(3):700–12. doi:10.1016/j.freeradbiomed.2011.05.024

39. Hu H, Takano N, Xiang L, Gilkes DM, Luo W, Semenza GL. Hypoxia-inducible factors enhance glutamate signaling in cancer cells. Oncotarget. Oct 15 2014;5(19):8853–68. doi:10.18632/oncotarget.2593

40. Wolfbeis OS. Luminescent sensing and imaging of oxygen: Fierce competition to the Clark electrode. 10.1002/bies.201500002. BioEssays. 2015/08/01 2015;37(8):921-928. 10.1002/bies.201500002

41. Wang X-d, Wolfbeis OS. Optical methods for sensing and imaging oxygen: materials, spectroscopies and applications. 10.1039/C4CS00039K. Chemical Society Reviews. 2014;43(10):3666-3761. doi:10.1039/C4CS00039K

42. Lee P, Chandel NS, Simon MC. Cellular adaptation to hypoxia through hypoxia inducible factors and beyond. Nature Reviews Molecular Cell Biology. 2020/05/01 2020;21(5):268-283. doi:10.1038/s41580-020-0227-y

43. Wagner BA, Venkataraman S, Buettner GR. The rate of oxygen utilization by cells. Free Radical Biology and Medicine. 2011/08/01/ 2011;51(3):700-712. 10.1016/j.freeradbiomed.2011.05.024

44. Kirchmair J, Göller AH, Lang D, et al. Predicting drug metabolism: experiment and/or computation? Nature Reviews Drug Discovery. 2015/06/01 2015;14(6):387-404. doi:10.1038/nrd4581

45. Nakazawa MS, Keith B, Simon MC. Oxygen availability and metabolic adaptations. Nat Rev Cancer. Sep 23 2016;16(10):663-73. doi:10.1038/nrc.2016.84

46. Sun RC, Denko NC. Hypoxic regulation of glutamine metabolism through HIF1 and SIAH2 supports lipid synthesis that is necessary for tumor growth. Cell Metab. Feb 4 2014;19(2):285–92. doi:10.1016/j.cmet.2013.11.022

47. Briggs KJ, Koivunen P, Cao S, et al. Paracrine Induction of HIF by Glutamate in Breast Cancer: EglN1 Senses Cysteine. Cell. Jun 30 2016;166(1):126–39. doi:10.1016/j.cell.2016.05.042

48. Koda S, Hu J, Ju X, et al. The role of glutamate receptors in the regulation of the tumor microenvironment. Review. Front Immunol. 2023-February-01 2023;14doi:10.3389/fimmu.2023.1123841

49. Sitkovsky MV, Hatfield S, Abbott R, Belikoff B, Lukashev D, Ohta A. Hostile, Hypoxia–A2- Adenosinergic Tumor Biology as the Next Barrier to Overcome for Tumor Immunologists. Cancer Immunology Research. 2014;2(7):598. doi:10.1158/2326-6066.CIR-14-0075

50. Hatfield SM, Kjaergaard J, Lukashev D, et al. Immunological mechanisms of the antitumor effects of supplemental oxygenation. Science Translational Medicine. 2015;7(277):277ra30. doi:10.1126/scitranslmed.aaa1260

51. Hardie DG, Schaffer BE, Brunet A. AMPK: An Energy-Sensing Pathway with Multiple Inputs and Outputs. Trends Cell Biol. Mar 2016;26(3):190–201. doi:10.1016/j.tcb.2015.10.013

52. Wortel IMN, van der Meer LT, Kilberg MS, van Leeuwen FN. Surviving Stress: Modulation of ATF4-Mediated Stress Responses in Normal and Malignant Cells. Trends Endocrinol Metab. Nov 2017;28(11):794–806. doi:10.1016/j.tem.2017.07.003

53. Derks J, Leduc A, Wallmann G, et al. Increasing the throughput of sensitive proteomics by plexDIA. Nature Biotechnology. 2023/01/01 2023;41(1):50-59. doi:10.1038/s41587-022-01389-w

54. Gordan JD, Thompson CB, Simon MC. HIF and c-Myc: sibling rivals for control of cancer cell metabolism and proliferation. Cancer Cell. Aug 2007;12(2):108–13. doi:10.1016/j.ccr.2007.07.006

55. Igarashi KJ, Kucinski I, Chan YY, et al. Physioxia improves the selectivity of hematopoietic stem cell expansion cultures. Blood Advances. 2023;7(14):3366–3377. doi:10.1182/bloodadvances.2023009668

56. Chen C, Tang Q, Zhang Y, Yu M, Jing W, Tian W. Physioxia: a more effective approach for culturing human adipose-derived stem cells for cell transplantation. Stem Cell Research & Therapy. 2018/05/24 2018;9(1):148. doi:10.1186/s13287-018-0891-4

57. Shin D-Y, Huang X, Gil C-H, Aljoufi A, Ropa J, Broxmeyer HE. Physioxia enhances T-cell development ex vivo from human hematopoietic stem and progenitor cells. Stem Cells. 2020;38(11):1454–1466. doi:10.1002/stem.3259

58. Wu MY, Yiang GT, Liao WT, et al. Current Mechanistic Concepts in Ischemia and Reperfusion Injury. Cellular Physiology and Biochemistry. 2018;46(4):1650–1667. doi:10.1159/000489241

59. Bugger H, Pfeil K. Mitochondrial ROS in myocardial ischemia reperfusion and remodeling. Biochim Biophys Acta Mol Basis Dis. Jul 1 2020;1866(7):165768. doi:10.1016/j.bbadis.2020.165768

60. Tiegs G, Horst AK. TNF in the liver: targeting a central player in inflammation. Semin Immunopathol. Jul 2022;44(4):445–459. doi:10.1007/s00281-022-00910-2

61. Hirao H, Nakamura K, Kupiec-Weglinski JW. Liver ischaemia-reperfusion injury: a new understanding of the role of innate immunity. Nat Rev Gastroenterol Hepatol. Apr 2022;19(4):239–256. doi:10.1038/s41575-021-00549-8

62. Zhou Z, Xu M-J, Gao B. Hepatocytes: a key cell type for innate immunity. Cellular & Molecular Immunology. 2016/05/01 2016;13(3):301-315. doi:10.1038/cmi.2015.97

63. Gan C, Cai Q, Tang C, Gao J. Inflammasomes and Pyroptosis of Liver Cells in Liver Fibrosis. Review. Front Immunol. 2022-May-30 2022;13doi:10.3389/fimmu.2022.896473

64. Pagé EL, Robitaille GA, Pouysségur J, Richard DE. Induction of Hypoxia-inducible Factor- 1&#x3b1; by Transcriptional and Translational Mechanisms *. Journal of Biological Chemistry. 2002;277(50):48403–48409. doi:10.1074/jbc.M209114200

65. Koshikawa N, Hayashi J, Nakagawara A, Takenaga K. Reactive oxygen species-generating mitochondrial DNA mutation up-regulates hypoxia-inducible factor-1alpha gene transcription via phosphatidylinositol 3-kinase-Akt/protein kinase C/histone deacetylase pathway. J Biol Chem. Nov 27 2009;284(48):33185–94. doi:10.1074/jbc.M109.054221

66. Jiang BH, Semenza GL, Bauer C, Marti HH. Hypoxia-inducible factor 1 levels vary exponentially over a physiologically relevant range of O_2_ tension. American Journal of Physiology-Cell Physiology. 1996/10/01 1996;271(4):C1172-C1180. doi:10.1152/ajpcell.1996.271.4.C1172

67. Rankin EB, Giaccia AJ. Hypoxic control of metastasis. Science. 2016;352(6282):175-180. doi:doi:10.1126/science.aaf4405

68. Mariotto AB, Etzioni R, Hurlbert M, Penberthy L, Mayer M. Estimation of the Number of Women Living with Metastatic Breast Cancer in the United States. *Cancer Epidemiology*, Biomarkers & Prevention. 2017;26(6):809–815. doi:10.1158/1055-9965.Epi-16-0889

69. Fuhrmann DC, Brüne B. Mitochondrial composition and function under the control of hypoxia. Redox Biology. 2017/08/01/ 2017;12:208-215. 10.1016/j.redox.2017.02.012

70. Vaupel P, Höckel M, Mayer A. Detection and characterization of tumor hypoxia using pO_2_ histography. Antioxid Redox Signal. Aug 2007;9(8):1221–35. doi:10.1089/ars.2007.1628

71. Posch W, Lass-Flörl C, Wilflingseder D. Generation of Human Monocyte-derived Dendritic Cells from Whole Blood. J Vis Exp. Dec 24 2016;(118)doi:10.3791/54968

72. Bolger AM, Lohse M, Usadel B. Trimmomatic: a flexible trimmer for Illumina sequence data. Bioinformatics. Aug 1 2014;30(15):2114–20. doi:10.1093/bioinformatics/btu170

73. Andrews S. FastQC. 2015.

74. Dobin A, Davis CA, Schlesinger F, et al. STAR: ultrafast universal RNA-seq aligner. Bioinformatics. 2012;29(1):15–21. doi:10.1093/bioinformatics/bts635

75. Liao Y, Smyth GK, Shi W. featureCounts: an efficient general purpose program for assigning sequence reads to genomic features. Bioinformatics. 2013;30(7):923–930. doi:10.1093/bioinformatics/btt656

76. Love MI, Huber W, Anders S. Moderated estimation of fold change and dispersion for RNA-seq data with DESeq2. Genome Biology. 2014/12/05 2014;15(12):550. doi:10.1186/s13059-014-0550-8

77. Subramanian A, Tamayo P, Mootha VK, et al. Gene set enrichment analysis: A knowledge-based approach for interpreting genome-wide expression profiles. Proceedings of the National Academy of Sciences. 2005;102(43):15545–15550. doi:doi:10.1073/pnas.0506580102

78. Yu G, Wang L-G, Han Y, He Q-Y. clusterProfiler: an R Package for Comparing Biological Themes Among Gene Clusters. OMICS: A Journal of Integrative Biology. 2012/05/01 2012;16(5):284-287. doi:10.1089/omi.2011.0118

79. Place TL, Domann FE, Case AJ. Limitations of oxygen delivery to cells in culture: An underappreciated problem in basic and translational research. Free Radical Biology and Medicine. 2017/12/01/ 2017;113:311-322.10.1016/j.freeradbiomed.2017.10.003

80. Sanford PA, Miller KG, Hoyt KO, Woolston BM. Deletion of biofilm synthesis in Eubacterium limosum ATCC 8486 improves handling and transformation efficiency. FEMS Microbiol Lett. Jan 17 2023;370doi:10.1093/femsle/fnad030

81. Woolston BM, Emerson DF, Currie DH, Stephanopoulos G. Rediverting carbon flux in Clostridium ljungdahlii using CRISPR interference (CRISPRi). Metab Eng. Jul 2018;48:243–253. doi:10.1016/j.ymben.2018.06.006

82. Colombani T, Eggermont LJ, Hatfield SM, et al. Oxygen-generating cryogels restore T cell-mediated cytotoxicity in hypoxic tumors. *bioRxiv*. 2020:2020.10.08.329805. doi:10.1101/2020.10.08.329805

83. Colombani T, Eggermont LJ, Rogers ZJ, et al. Biomaterials and Oxygen Join Forces to Shape the Immune Response and Boost COVID-19 Vaccines. Advanced Science. 2021/09/01 2021;8(18):2100316. 10.1002/advs.202100316

84. Specht H, Slavov N. Optimizing Accuracy and Depth of Protein Quantification in Experiments Using Isobaric Carriers. Journal of Proteome Research. 2021/01/01 2021;20(1):880- 887. doi:10.1021/acs.jproteome.0c00675

85. Specht H, Emmott E, Petelski AA, et al. Single-cell proteomic and transcriptomic analysis of macrophage heterogeneity using SCoPE2. Genome Biol. Jan 27 2021;22(1):50. doi:10.1186/s13059-021-02267-5

86. Petelski AA, Emmott E, Leduc A, et al. Multiplexed single-cell proteomics using SCoPE2. Nature Protocols. 2021/12/01 2021;16(12):5398-5425. doi:10.1038/s41596-021-00616-z

87. Demichev V, Messner CB, Vernardis SI, Lilley KS, Ralser M. DIA-NN: neural networks and interference correction enable deep proteome coverage in high throughput. Nature Methods. 2020/01/01 2020;17(1):41-44. doi:10.1038/s41592-019-0638-x

88. Leek JT, Johnson WE, Parker HS, Jaffe AE, Storey JD. The sva package for removing batch effects and other unwanted variation in high-throughput experiments. Bioinformatics. Mar 15 2012;28(6):882–3. doi:10.1093/bioinformatics/bts034

89. Ritchie ME, Phipson B, Wu D, et al. limma powers differential expression analyses for RNA-sequencing and microarray studies. Nucleic Acids Res. Apr 20 2015;43(7):e47. doi:10.1093/nar/gkv007

